# The mammalian-adaptive PB2-E627K substitution preserves viral fitness of clade 2.3.4.4b H5N1 HPAIV in birds

**DOI:** 10.64898/2026.10.06.757006

**Authors:** Elizabeth Billington, Sofia Riccio, Maryn D. Brown, Benjamin C. Mollett, Jessica L. Quantrill, Simon Johnson, Cecilia Di Genova, Jiayun Yang, Jean-Remy Sadeyen, Audra-Lynne Schlachter, Kelly J Roper, Jafar Hassan, Benjamin Clifton, Kajal Ralh, Caroline J. Warren, Georgina Ward, Dilhani de Silva, Alejandro Nunez, Holly A. Coombes, Marek J. Slomka, Munir Iqbal, Wendy S. Barclay, Thomas P. Peacock, Ashley C. Banyard, Joe James

## Abstract

Since emerging in 2020, clade 2.3.4.4b H5N1 high pathogenicity avian influenza virus (HPAIV) has disseminated globally, causing substantial infection and mortality in wild birds and poultry. Additionally, numerous spill-over events into mammals have occurred, including mass mortalities and sporadic human infections. Such events can drive the acquisition of mammalian-adaptive mutations, which further increase the risk to humans; the most well characterised mutation is the glutamate to lysine change at amino acid position 627 in the polymerase basic 2 protein (PB2-627K). Across avian influenza viruses, PB2-627K enhances polymerase activity and replication in mammalian cells, increasing zoonotic and pandemic potential. However, bioinformatic analyses indicate that PB2-627K remains rare among clade 2.3.4.4b H5N1 viruses circulating in avian populations. Here, we investigated the impact of PB2-627K on viral fitness in avian hosts to assess the potential for mammalian-adapted viruses to re-establish in birds, mimicking infection of avian species following scavenging on mammalian species where PB2-627K has already emerged. Using a representative H5N1 virus (European genotype AB), PB2-627K increased polymerase activity and replication in human cells without compromising replication in avian cells. A mammalian-derived PB2-627K H5N1 isolate from a fox exhibited comparable replication kinetics and transmission efficiency in chickens and ducks to a closely related PB2-627E virus. Notably, PB2-627K remained genetically stable at consensus level following infection and transmission in both avian hosts. These findings demonstrate that mammalian-adaptive mutations acquired during spill-over can be maintained in avian populations, potentially elevating zoonotic risk and underscoring the importance of surveillance for such mutations in birds.

**Importance:** Clade 2.3.4.4b H5N1 high pathogenicity avian influenza viruses (HPAIVs) have caused widespread outbreaks in birds and increasingly frequent spill-over into wild and domestic mammals, including recent infections in cattle. Greater circulation in mammalian hosts raises concern that viruses may acquire mutations associated with human adaptation. We assessed the risk of such “pre-adapted” viruses, already possessing mammalian-adaptive markers, re-entering and persisting in avian populations. We show that the key adaptation PB2-627K does not compromise viral fitness in birds and remains genetically stable following transmission in chickens and ducks. These findings indicate that mammalian-adaptive mutations acquired during spill-over can be maintained in avian reservoirs, potentially lowering the barrier to zoonotic infection. This work highlights the importance of genomic surveillance in both mammalian and avian populations to support any alterations to zoonotic risk.

## INTRODUCTION

In 1996, a lineage of H5 high pathogenicity avian influenza virus (HPAIV) was detected in Guangdong, China, termed Goose/Guangdong (gs/Gd) (1). Since its emergence, this lineage has disseminated globally, largely mediated by wild bird migration, and has diversified through mutation and reassortment into multiple clades (based on the HA gene) and genotypes respectively (2). In 2020, a H5N1 HPAIV belonging to clade 2.3.4.4b was detected in wild birds and poultry across Europe. The virus subsequently expanded and became widespread in wild bird populations during 2021, before transatlantic spread to North America in late 2021 to early 2022 (3, 4), then to South America in late 2022 (5) and, for the first time, to the Antarctic region in late 2023 (6), where it has spread along island populations in the Atlantic, Southern, and Indian Oceans (7) to reach Australia in June 2026 (8).

The current H5N1 panzootic has caused devastating losses in poultry and unprecedented mortality in wild bird populations, particularly among waterfowl (Anseriformes) and gulls (Charadriiformes) (5, 9). In addition, numerous infections have been documented in wild mammals worldwide (10, 11), most commonly arising from predation or scavenging of infected birds (10, 11). Potential instances of sustained wild mammal-to-mammal transmission have been suggested, such as during mass die-offs of marine mammals in South America and the Antarctic region (7, 10, 12–14). Beyond wildlife, infections have also been reported in captive mammal species in domestic (15), zoological (16), and farm settings (10, 17, 18); most notable is the widespread outbreak in dairy cattle in the United States of America (USA) following the first reported case on 25 March 2024 (10, 18). Sporadic human cases have also occurred, with 92 confirmed human infections, three of which have been fatal, with clade 2.3.4.4b H5N1 HPAIV up until March 2026 (18).

Cross-species transmission of avian influenza viruses (AIVs) is often accompanied by adaptive genetic changes across the viral genome, enabling the virus to overcome host-specific molecular, physiological, or environmental barriers to infection, replication, and onward transmission (19, 20). Such adaptations are thought to occur in a stepwise fashion, reflecting both the nature of the host barrier and the ease with which it can be overcome (19, 20). In mammalian hosts, the earliest adaptive changes frequently occur in the viral RNA-dependent RNA polymerase (vRDRP) complex, enhancing polymerase activity and viral replication kinetics in the new host species (19–22). Additional mutations may subsequently arise in other regions of the genome, enabling improved receptor utilisation, evasion of host innate immunity, or enhanced environmental stability (e.g., altered pH threshold of fusion improving aerosol transmission) (19, 20).

One of the most well-characterised mammalian-adaptive mutations in AIVs is the substitution of glutamate (E) with lysine (K) at position 627 of the polymerase basic protein 2 (PB2-E627K) (23–27). PB2 forms part of the vRDRP complex, together with PB1 and PA, and acquisition of PB2-627K markedly enhances polymerase activity and viral replication in mammalian cells (24, 26). This phenotype is mediated through improved utilisation of mammalian ANP32 proteins, key host cofactors required for influenza virus replication (26). Whereas avian-adapted polymerases efficiently interact with avian ANP32A, they are restricted in mammalian cells, a barrier that can be overcome by the PB2-627K substitution (28–31).

AIVs that possess PB2-627K, or other mammalian-adaptive mutations, therefore pose a greater zoonotic threat than their fully avian adapted counterparts. This may be through enhanced infectivity in humans, or by facilitating the establishment of sustained lineages in mammalian hosts, where further adaptation may occur. Although infection of wild mammals heightens zoonotic potential, the risk of subsequent human exposure is typically low, due to infrequent direct encounters with dead or clinically affected wild mammals (32). Close human interaction with captive poultry species still represents the greatest risk for zoonotic infection (19, 32). Consequently, a key concern during the ongoing H5N1 panzootic is whether mammal-adapted viruses can ‘spill back’ into avian populations, potentially reintroducing mammalian-adaptive mutations into wild bird reservoirs and onwards to poultry. Such events could significantly increase zoonotic risk by disseminating viruses with an enhanced ability to infect mammals.

The role of PB2-627K in the mammalian adaptation of clade 2.3.4.4b H5N1 is underscored by its frequent detection in natural and experimental mammalian infections compared to avian counterparts (11, 27, 33–35). In addition, a conversion rate from E to K at position 627 in PB2 of up to 40% has been observed across all gs/Gd lineage H5 HPAIVs following infection of humans (36). By comparison, the vast majority of avian isolates retain PB2-627E (27). Owing to diversity among ANP32 proteins, a limited number of avian species, such as ratites (e.g., ostriches and emus), have been demonstrated to select for viruses harbouring PB2-627K following infection (26, 37). However, there is no empirical evidence to suggest that wild aquatic birds, as the principal clade 2.3.4.4b reservoir hosts, or gallinaceous poultry (e.g., chickens and turkeys), which represent the main farmed species, exert similar selective pressures to select PB2-627K.

The apparent absence of PB2-627K among H5N1 clade 2.3.4.4b HPAIV sequences in wild birds during the current epizootic suggests two possible explanations; (**i**) infections of wild mammals represent epidemiological dead-ends with negligible onward transmission potential to wild bird reservoir species, or (**ii**) PB2-627K imposes a fitness cost in avian hosts, leading to reversion or competitive exclusion by PB2-627E. The magnitude of such a fitness cost remains poorly characterised yet appears to vary between viral lineages (38).

In this study, we investigated the fitness consequences of the PB2-627K mutation in clade 2.3.4.4b H5N1 HPAIV (genotype AB) in avian hosts. We examined the effects of PB2-627K on polymerase activity and viral replication in both mammalian and avian cells. We compared replication kinetics, disease progression, transmission dynamics, and genetic stability *in vivo*, in ducks and chickens, using two H5N1 HPAIVs representative of the predominant AB genotype circulating in the United Kingdom (UK) and Europe since 2022. These viruses represented an avian-adapted variant (possessing PB2-627E) and a highly closely related mammalian-adapted PB2-627K containing variant that had naturally acquired this polymorphism.

## RESULTS

### PB2-627K is enriched in mammalian-derived H5 clade 2.3.4.4b HPAIV sequences but rare in most avian hosts

From the full dataset of avian-derived clade 2.3.4.4b H5 HPAIV PB2 sequences (*n* = 27,813) 98.9% (n=27,526) of the sequences encoded 627E compared to the 1.1% (n=287) encoding 627K from 2002-2025 (Fig. 1A). This pattern was consistent across the well sampled orders including Anseriformes (waterfowl such as ducks, geese, and swans) (99.7% (n=10,103) E, 0.3% K (n=27)), Galliformes (poultry such as chickens and turkeys) (98.6% E (n=8425), 1.4% K (n=117)) and Charadriiformes (shorebirds, gulls, and waders) (98.6% E (n=2935), 1.4% K (n=40)), where 627K represented only a minor fraction of the sequences (Fig. 1A). Similar trends were observed across other orders such as Accipitriformes (birds of prey) (98.5% E (n=2274), 1.5% K (n=35)), Passeriformes (perching birds) (97.0% E (n=820), 3.0% K (n=25)), and Strigiformes (owls) (99.5% E (n=822), 0.5% K (n=4)), reinforcing the strong conservation of 627E in avian hosts. In contrast, a higher relative frequency of 627K was observed in ratite orders, including Casuariiformes (large flightless birds) (48.0% E (n=12), 52.0% K (n=13)) and Rheiformes (rheas) (16.7% E (n=2), 83.3% K (n=10)), although these groups were represented by small sample sizes (Fig. 1A).

**Figure 1.**
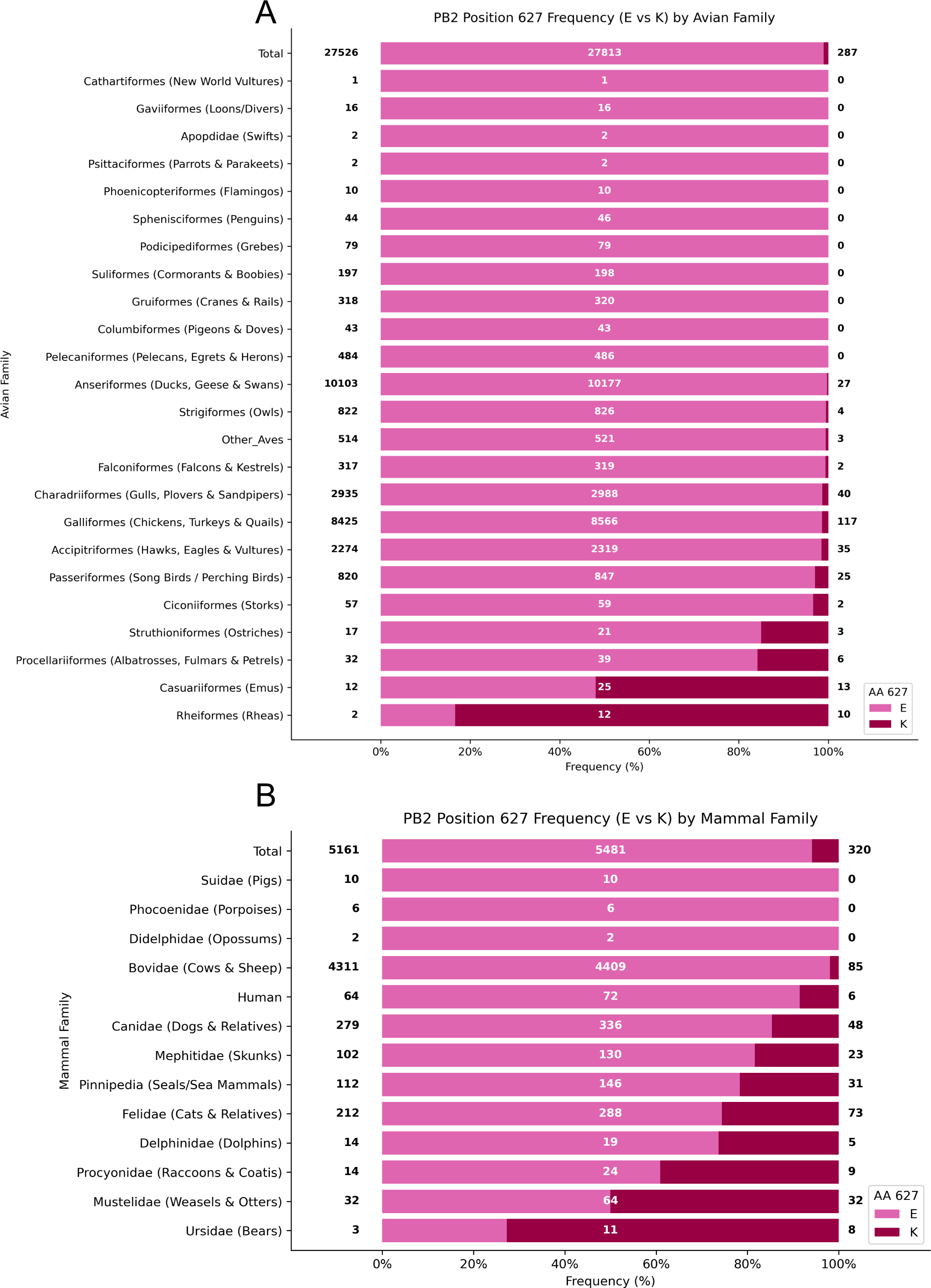
Frequency of PB2 residue 627 variants in viral sequences derived from different avian and mammalian families. (**A**) The Relative frequency (%) of amino acids E (glutamic acid) and K (lysine) at PB2 position 627 in H5 Clade 2.3.4.4b HPAIV sequences, grouped by avian family/order. Within each family, bars are normalized to 100% and divided into E (light pink) and K (dark maroon). Annotated numbers indicate sequence counts: the left value denotes the number of sequences with E, the right value denotes the number with K, and the central value gives the total number of sequences for that family. The “Total” row summarizes the overall distribution of E versus K across all avian families included. PB2 sequences were from H5 Clade 2.3.4.4b HPAIVs obtained during the period 2002-2025. (**B**) The Relative frequency (%) of amino acids E (glutamic acid) and K (lysine) at PB2 position 627 in influenza A virus sequences, grouped by mammalian family/order. Within each family, bars are normalized to 100% and divided into E (light pink) and K (dark maroon). Annotated numbering is as for panel **A**.

For the mammalian-derived clade 2.3.4.4b H5 HPAIV PB2 sequences (n = 5,481), position 627 was still predominantly E, although K was observed at a higher frequency than in avian hosts. Across the full dataset of mammalian derived sequences, 627E accounted for 94.2% (n=5161) of sequences, compared to 5.8% for 627K (n=320) (Fig. 1B). However, this overall distribution varied substantially between mammalian families. In Bovidae (cattle and related ruminants), which comprised the majority of sequences, 627E was highly predominant (98.1% E (n=4311), 1.9% K (n=85)) (Fig. 1B). Similarly, Suidae (pigs), Phocoenidae (porpoises), and Didelphidae (opossums) showed complete representation of 627E, although these groups were represented by small sample sizes.

Higher proportions of 627K were observed in several other mammalian families. For example, 627K was present in Human (8.3% (n=6)), Canidae (dogs and foxes) (14.3% (n=48)), Felidae (cats) (26.4% (n=73)), Mephitidae (skunks) (20.8% (n=23)), and Pinnipedia (seals and sea lions) (23.3% (n=31)), with similar levels in Delphinidae (dolphins) (26.3% (n=5)). In Mustelidae (weasels, mink, and related species), 627K accounted for 50.0% (n=32) of sequences, while Ursidae (bears) showed 72.7% (n=8) 627K, although both were based on limited numbers of sequences (Fig. 1B).

Overall, these data demonstrate that PB2-627E remains strongly conserved in avian clade 2.3.4.4b H5 HPAIVs, whereas PB2-627K is enriched in mammalian-derived viruses and reaches particularly high frequencies in several carnivore families.

### The PB2-E627K polymorphism does not impact the replication kinetics of clade 2.3.4.4b H5N1 HPAIV in avian cells

We next examined the effect of the PB2-E627K polymorphism on influenza virus polymerase activity in avian and human cells. A polymerase minigenome assay was established using the polymerase complex of a genotype AB clade 2.3.4.4b H5N1 HPAIV, representative of a dominant European genotype during the clade 2.3.4.4b epizootic. The PB2 was mutated, creating two PB2 variants, one possessing PB2-627E (avian), and one possessing PB2-627K (mammalian-adaptive). The relative activities were compared, with polymerase activity normalised to a PB2-deficient control (ΔPB2) (Fig. 2A). In human 293T cells, H5N1 genotype AB polymerase complexes carrying PB2-627E showed very low activity (Fig. 2A). However, complexes possessing PB2-627K exhibited a highly statistically significant increase in activity compared to the PB2-627E comparator (Fig. 2A). In avian cell lines (chicken, DF-1 and CLEC; duck, CCL-141), the PB2-627E baseline polymerase activity was higher compared to that in 293T cells. Possession of PB2-627K yielded a small increase in activity, which was only statistically significant in the duck cells (Fig. 2A).

**Figure 2.**
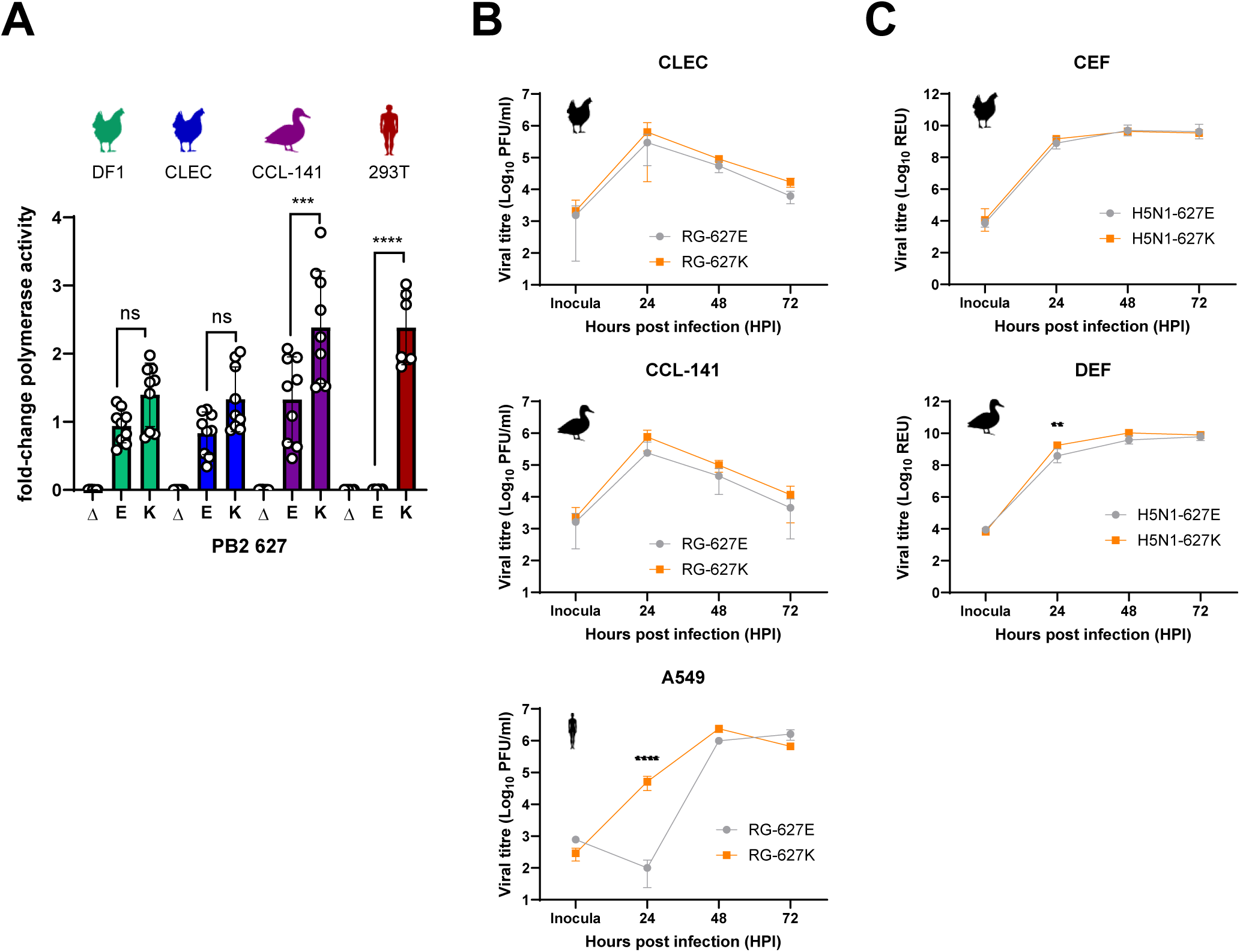
Impact of the PB2-E627K polymorphism on polymerase activity and viral replication in avian and human cells. **(A)** Fold-change in polymerase activity measured by mini-replicon assay using PB1, PB2, PA, and NP gene segments from a genotype AB Clade 2.3.4.4b H5N1 HPAIV. Constructs differed only at PB2 residue 627 (E or K) or lacked PB2 (Δ). Assays were conducted in chicken (DF-1 and CLEC), duck (CCL-141) and human (293T) cell lines. (**B**) Replication kinetics of reverse genetics viruses differing only at PB2 position 627 (RG-627E and RG-627K) in chicken (CLEC), duck (CCL-141) and human (A549) cells. Both viruses contained the six internal gene segments (PB2, PB1, PA, NP, M, NS) from a genotype AB Clade 2.3.4.4b H5N1 HPAIV and the HA and NA segments from A/Puerto Rico/8/1934 (H1N1). Supernatants titrated on MDCK cells by plaque assay. (**C**) Replication kinetics of two naturally occurring Clade 2.3.4.4b H5N1 HPAIVs that are highly similar but differ at PB2 residue 627 (E or K) (H5N1-627E and H5N1-627K) in primary chicken (CEF) and duck (DEF) fibroblasts. Viral titre was measured by H5 RT-PCR of RNA extracted from supernatants. (**A-C**) Data was compared for significance in GraphPad Prism v10 using two-way ANOVA, unless indicated with comparisons were not statistically significant (ns = ≥ P 0.05, ** = P < 0.01, *** = P < 0.001, **** = P < 0.0001).

We next assessed the impact of PB2-627 polymorphisms on the viral replication kinetics, using RG viruses which differed only in the possession of PB2-627E (RG-627E) or PB2-627K (RG-627K). Consistent with the polymerase assay data, in human A549 cells, RG-627K showed statistically significantly higher titres than RG-627E at 24 hpi, however titres were comparable by 72 hpi (Fig. 2B). In both duck and chicken cells, RG-627K showed slightly higher, yet not statistically significant, viral replication at all timepoints compared to RG-627E (Fig. 2B). We next investigated replication kinetics in wild type clade 2.3.4.4b H5N1 genotype AB viruses, H5N1-627E or H5N1-627K (Fig. 2C). Both viruses replicated equally well in CEF and DEF, with no statistically significant difference in replication observed apart from a modest, yet statistically significant increase in replication of PB2-627K at 24 hpi in duck cells (DEFs) (Fig. 2C).

### PB2-E627K does not negatively impact infectivity, replication or pathogenicity of clade 2.3.4.4b H5N1 HPAIV in ducks or chickens

We next investigated the impact of PB2-627 in both chickens and ducks on infection kinetics. Separate groups of ducks, representing waterfowl as the natural reservoir of genotype AB H5N1, and chickens, the primary commercially relevant poultry species, were inoculated with either H5N1-627E or H5N1-627K (Fig. S1).

Both H5N1-627E and H5N1-627K caused 100% infection in donor (D0) ducks, with all individuals shedding vRNA from the oropharyngeal (OP) and cloaca (C) by 3 days post infection (dpi) (Fig. 3A and B). For both PB2-627 variants, mean OP vRNA shedding peaked at 3 dpi with positive detections persisting until 6 dpi, when mortality reached 100% in the H5N1-627E group, and the single remaining duck in the H5N1-627K group ceased shedding (Fig. 3A, D). Cloacal shedding was generally lower and more variable than OP shedding in both groups; however, all ducks shed vRNA from the cloaca at least once during the study (Fig. 3B). Peak vRNA levels were reached earlier in swabs collected from ducks infected with H5N1-627E (2-3 dpi) than those infected with H5N1-627K (4-6 dpi) (Fig. 3B). The total mean OP and C shedding, calculated as the area under the curve (AUC), was not significantly different between the H5N1-627E and H5N1-627K groups (Fig. 3C). Similar levels of vRNA were detected in the environment (drinking and pond waters) of both H5N1-627E and H5N1-627K infected ducks, with environment vRNA levels closely following the period that ducks were actively shedding vRNA (Fig. S2A-B, Fig 3A-B).

**Figure 3.**
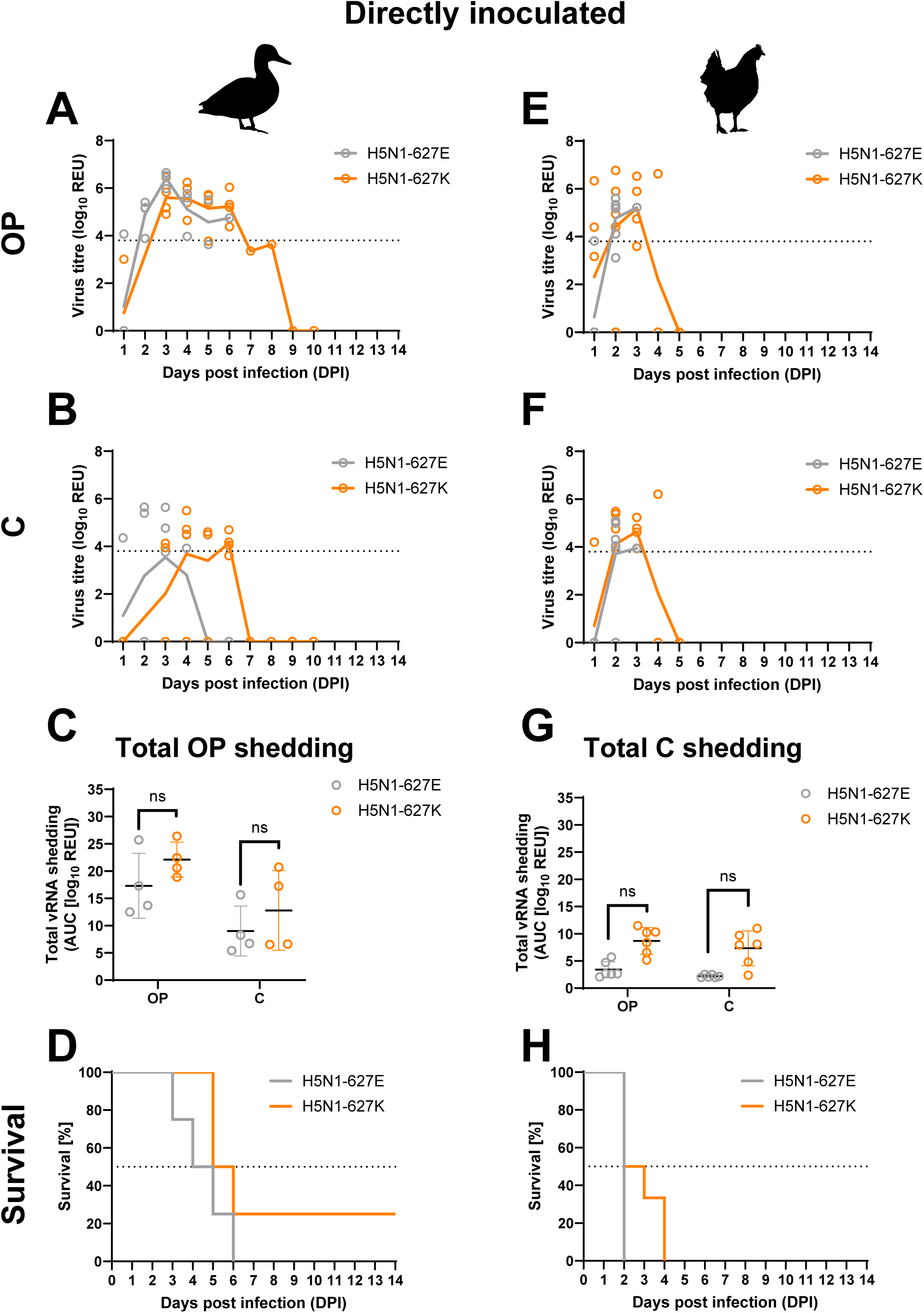
Viral shedding and mortality in chickens and ducks infected with near-identical clade 2.3.4.4b H5N1 HPAIVs differing at PB2-627. Ducks and chickens were inoculated via the intraocular–intranasal route with 10^4^ EID_50_ of clade 2.3.4.4b H5N1 viruses that are nearly genetically identical, except for the naturally occurring amino acid at PB2-627, E (H5N1-627E) or K (H5N1-627K). **A-F**, Viral RNA shedding kinetics measured by RT-PCR of RNA extracted from oropharyngeal (OP; A, E) and cloacal (C; B, F) swabs collected from inoculated ducks (**A-B**) and chickens (**E-F**). **C** and **G**, Area under the curve (AUC) analysis of individual birds, showing total mean shedding in ducks (**C**) and chickens (**G**). **D, H,** Survival curves for ducks (**D**) and chickens (**H**) following inoculation.

Mortality in ducks was similar between groups, reaching 100% for H5N1-627E and 75% for H5N1-627K (Fig. 3D). The progression of mortality also followed a comparable trend, with ducks infected with H5N1-627E exhibiting more rapid clinical disease progression than those infected with H5N1-627K with median survival of 4.5 days compared to 6 days respectively, although this difference was not statistically significant (Fig. 3D, Table S1, Table S4). The single surviving H5N1-627K infected duck was also serologically positive against H5 ELISA and Hemagglutination inhibition (HI) (Fig S3).

In chickens, both H5N1-627E and H5N1-627K also caused 100% infection, with all chickens shedding vRNA from the OP or C (Fig. 3E-F). vRNA was first detected at the first sampling point (1 dpi) and continued until all chickens died (Fig. 3E, F and H). Cloacal shedding followed a similar temporal pattern to OP shedding but occurred at lower titres (Fig. 3F). Chickens inoculated with H5N1-627K exhibited higher total mean OP and C shedding, but this was not statistically significant (Fig. 3G). Mortality reached 100% in both groups, with clinical disease leading to mortality progressing slightly faster for H5N1-627E (median survival 2 dpi, 100% mortality by 2 dpi) than for H5N1-627K (median survival 3 days, 100% mortality by 4 dpi), although this difference was not statistically significant (Fig. 3H, Table S1).

Collectively, these findings indicate that PB2-627K does not reduce infectivity, shedding, or virulence of clade 2.3.4.4b H5N1 HPAIV in either ducks or chickens.

### The PB2-E627K substitution does not reduce transmission efficiency of clade 2.3.4.4b H5N1 HPAIV in ducks or chickens

We also investigated if PB2-627K had any impact on intra-species transmission efficiency among ducks and chickens by introducing naïve contact ducks or chickens to the infected groups at 1 dpi, and co-housed these recipient (R1) birds for the duration of the study.

All contact ducks became infected, corresponding to a transmission efficiency of 100% (8/8) for both H5N1-627E and H5N1-627K (Fig. 4A-B, Table S1). Transmission occurred earlier with H5N1-627E, with all R1 ducks (8/8) shedding vRNA from OP or C cavities by 3 dpi (2 days post-contact [dpc]), compared with 75% (6/8) of R1 ducks shedding H5N1-627K vRNA at the same time point (Fig. 4A-B). By 4 dpi (3 dpc), all ducks were shedding vRNA from the OP and/or C cavity. In the H5N1-627E group, shedding ceased as all R1 ducks succumbed to infection, whereas in the H5N1-627K group, shedding gradually declined as two R1 ducks cleared the infection (Fig. 4A-B, D). This resulted in a mean interval from first contact to first vRNA-positive detection of 1.88 days (SD ± 0.35) for H5N1-627E and 2.25 days (SD ± 0.46) for H5N1-627K (Table S1). No significant difference was observed in total vRNA shedding between the H5N1-627E and –627K groups from either the OP or C (Fig. 4C). Mortality did not differ significantly between groups, with mortality in H5N1-627E first being observed at 5 dpi (4 dpc), reaching 100% (8/8) in contact ducks by 6 dpi (5 dpc). For H5N1-627K mortalities reached 75% (6/8) in H5N1-627K contact ducks between 5 dpi (4 dpc) and 7 dpi (6 dpc) (Fig. 4D). The two surviving R1 ducks in the H5N1-627K group also seroconverted to H5N1, confirming infection in these individuals (Fig. S3).

**Figure 4.**
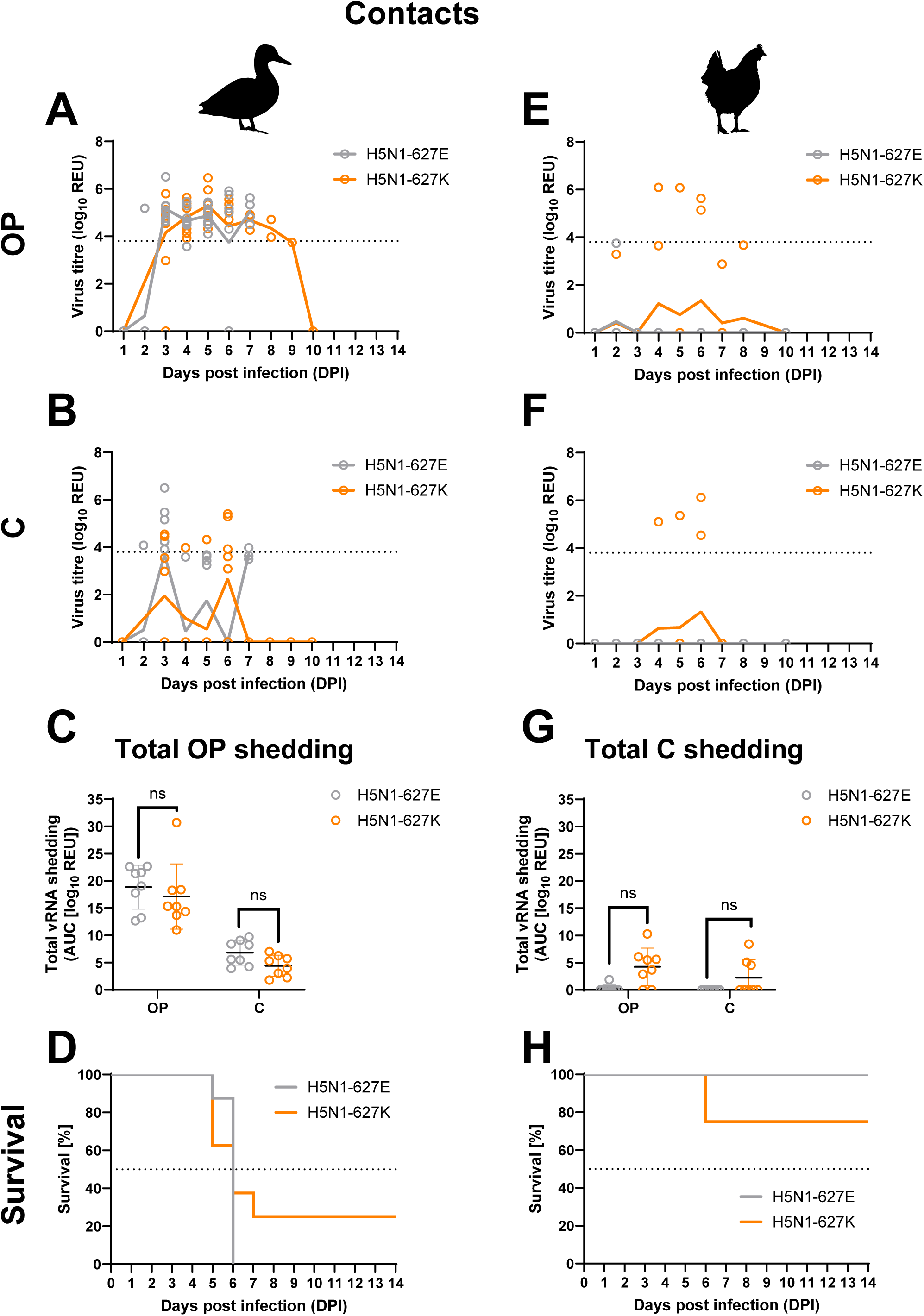
Intraspecies transmission dynamics of clade 2.3.4.4b H5N1 HPAIVs carrying PB2-627E or PB2-627K among chickens and ducks. Ducks and chickens were co-housed, at 1-day post-infection (dpi), with ducks and chickens which were inoculated with 2.3.4.4b H5N1 viruses that are nearly genetically identical, except for the naturally occurring amino acid at PB2-627, E (H5N1-627E) or K (H5N1-627K). **A-F**, Viral RNA shedding kinetics measured by RT-PCR of RNA extracted from oropharyngeal (OP; A, E) and cloacal (C; B, F) swabs collected from contact ducks (**A-B**) and chickens (**E-F**). **C** and **G**, Area under the curve (AUC) analysis of individual birds, showing total mean shedding in ducks (**C**) and chickens (**G**). **D** and **H,** Survival curves for contact ducks (**D**) and chickens (**H**) following inoculation.

In chickens, inefficient transmission was seen for both viruses, with H5N1-627E failing to transmit to any R1s, as evidenced by the absence of clinical signs, detectable vRNA shedding, and seroconversion (Fig. 4E-H and Fig S3). Consequently, the transmission efficiency of H5N1-627E was 0% (0/8). In contrast, H5N1-627K transmitted to two contact chickens, corresponding to a transmission efficiency of 25% (2/8) (Fig. 4E-F). In the chickens where transmission occurred, H5N1-627K vRNA was detected in both OP and C swabs from 3541 at 5 and 6 dpi (4 and 5 dpc), and from 3538 at 6 dpi (5 dpc) (Fig. 4E-F). R1 chickens 3541 and 3538 succumbed to infection at 6 and 7 dpi respectively, (Fig. 4H). vRNA was also detected in the OP and C swabs of chicken 3539 at 4 dpi (3 dpc), however this bird did not display any clinical signs or seroconvert to homologous antigen and was therefore not considered robustly infected. None of the remaining five R1 chickens in the H5N1-627K group displayed any clinical signs or seroconverted to the homologous antigen (Fig. S3). The mean interval from first contact to the onset of vRNA shedding in positive contact birds was 3.5 days (SD ± 0.71). All infected H5N1-627K R1s exhibited short shedding durations, as they succumbed to infection within 48 hours of the first vRNA-positive swab.

### PB2-E627K does not alter tissue tropism, viral distribution, or pathology of clade 2.3.4.4b H5N1 HPAIV in ducks or chickens

In ducks, vRNA and viral NP antigen were detected in multiple tissues from birds that succumbed to infection and were taken for opportunistic postmortem examination, indicating widespread systemic dissemination (Fig. 5A and C). vRNA was present across respiratory, gastrointestinal, lymphoid and parenchymal tissues, with the highest titres generally observed in the brain. Consistent with the vRNA data, the most intense NP immunolabelling was detected in the cerebrum, with more limited staining observed in other tissues, including the liver, pancreas, trachea and syrinx. Although direct comparisons were constrained by differences in sampling time and clinical outcome, no clear differences in tissue distribution or antigen staining were evident between H5N1-627E- and H5N1-627K-infected ducks. Both D0 and R1 ducks developed systemic infection and exhibited similar patterns of viral distribution in tissues. In contrast, surviving H5N1-627K contact ducks sampled at 14 dpi showed no detectable NP immunolabelling and only limited residual vRNA, consistent with substantial clearance of infection.

**Figure 5.**
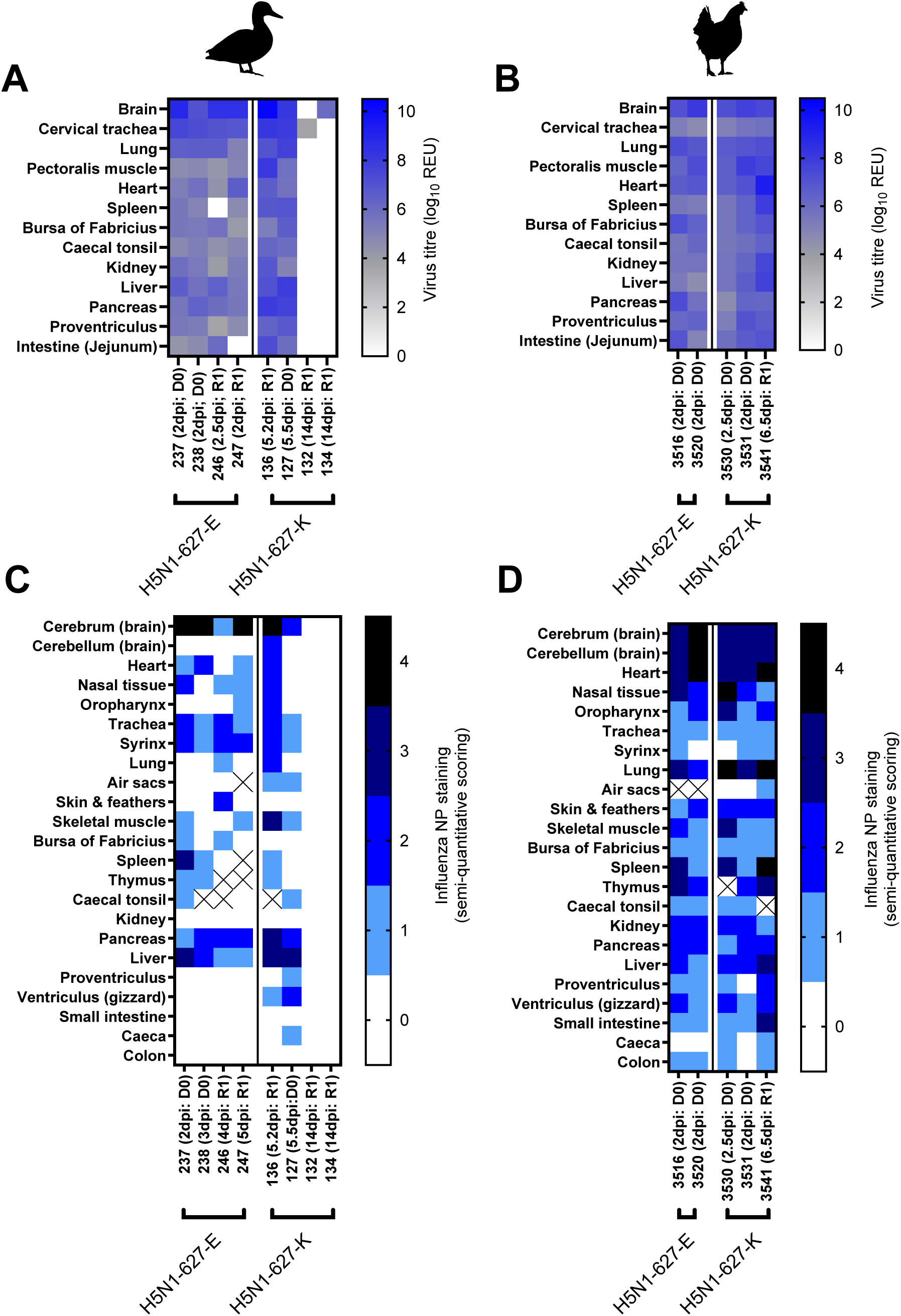
Viral tissue tropism and immunohistochemical detection of clade 2.3.4.4b H5N1 HPAIVs carrying PB2-627E or PB2-627K in chickens and ducks. Tissues representing various organ systems, including the nervous, respiratory, cardiovascular, lymphoid, digestive, and musculoskeletal systems, were collected from ducks (**A, C**) and chickens (**B, D**) were inoculated with 10^4^ EID_50_ of clade 2.3.4.4b H5N1 viruses carrying either PB2-627E (H5N1-627E) or PB2-627K (H5N1-627K). Viral RNA loads and antigen distribution were assessed in multiple tissues obtained during post-mortem examination (PME) following mortality. **A** and **B**, heatmaps depicting viral RNA levels, measured by RT-PCR and expressed as relative equivalency units (REU), in organs from inoculated chickens (**A**) and ducks (**B**). Darker shades of blue correspond to higher viral RNA titres, shades of grey indicate sub-threshold results (>Ct 36), and white indicates negative sample. **C** and **D**. Immunohistochemical (IHC) scoring of viral antigen in the corresponding tissues from chickens (**C**) and ducks (**D**). IHC scores indicate antigen abundance (0 = none, 4 = abundant). “X” denotes samples that could not be collected or tested.

Chickens infected with either H5N1-627E or H5N1-627K exhibited extensive viral dissemination, with high vRNA titres detected throughout the respiratory, lymphoid, gastrointestinal and parenchymal tissues examined (Fig. 5B). vRNA titres were broadly comparable between virus variants, with no evidence of altered tissue tropism associated with the PB2 E627K substitution. Similar patterns of tissue distribution were observed in directly infected and contact birds. Consistent with these findings, NP antigen was widely distributed across multiple organs (Fig. 5D). The most intense and consistent immunolabelling was observed in the cerebrum, cerebellum and heart, where staining was uniformly moderate to marked across all birds. Antigen was also frequently detected in respiratory tissues, spleen, liver, pancreas, kidney and skeletal muscle, demonstrating widespread tissue involvement irrespective of virus variant.

Overall, the PB2-E627K substitution did not measurably alter tissue tropism, viral dissemination, or pathological distribution in either ducks or chickens.

### In both ducks and chickens, the PB2-E627K polymorphism is maintained without reversion during infection and transmission of clade 2.3.4.4b H5N1 HPAIV

Whole-genome sequencing using Oxford Nanopore Technology was performed on selected clinical samples (swabs and tissues) from H5N1-627K infected chickens and ducks. Samples were chosen to, (i) maximise representation of viral sequences from each infected bird; (ii) capture the virus at the latest detectable stage of infection; and (iii) ensure sufficient vRNA was present to enable successful sequencing (Fig. 6). From H5N1-627K infected chickens, only 3/8 R1 chickens shed vRNA, and of those three only sporadic or single day shedding was detected. Due to lower vRNA titres of C swabs, a smaller subset of chickens was sequenced from both cavities. More duck samples were available to be sequenced due to a higher transmission rate and the longer shedding window (Fig. 6). Tissue samples from D0 and R1 ducks and chickens were also sequenced, to assess whether reversion had occurred to PB2-627E. All samples sequenced maintained 627K at consensus level, with relatively few additional mutations were detected (Table S2).

**Figure 6.**
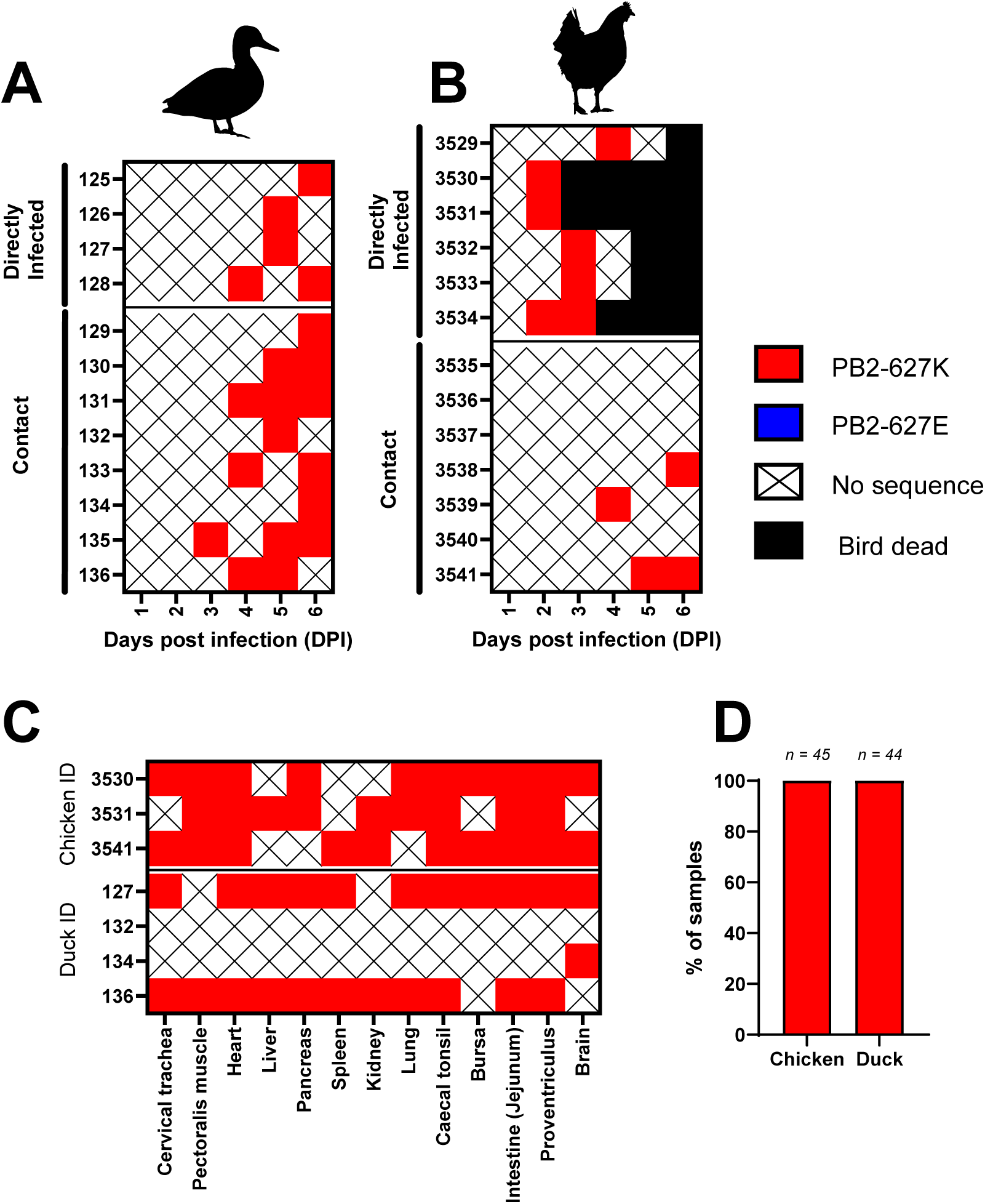
Genetic stability of 627K in PB2 in chickens and ducks. Swab (**A** and **B**) and tissue (**C**) samples selected for whole genome sequencing (WGS) to assess the genetic stability of PB2-627. Samples were collected from chickens and ducks either inoculated with clade 2.3.4.4b H5N1 virus with the amino acid lysine (K) at position 627 in the PB2 protein (H5N1-627K), or from contact birds. Sequencing results indicate the presence of PB2-627E (blue) or PB2-627K (red) at the consensus level, with total percentage of E or K in chicken or duck samples shown in **D**.

Only a small number of additional amino acid polymorphisms were identified, most frequently in duck swabs from 5-6 dpi (Table S2). Most were unique to individual animals, and no mutation was observed repeatedly across multiple ducks or chickens. Most substitutions occurred within polymerase genes, particularly PB2, which accounted for seven of the 13 amino acid changes detected. Four of these seven (V553I, T666Q, A627G and G673D) occurred in the PB2 627 domain, however little is reported on these mutations within the literature.

Collectively, these data indicate that PB2-627K is genetically stable during infection and transmission in both ducks and chickens, consistent with the absence of a detectable fitness cost in either host species.

## DISCUSSION

The global expansion of clade 2.3.4.4b H5N1 HPAIV since 2020 has profoundly altered the ecology and evolutionary dynamics of HPAIV. The ongoing panzootic has resulted in unprecedented levels of infection and mortality in wild bird populations, widespread outbreaks in poultry, and an unparalleled number of spillover infections into mammalian hosts, including evidence of sustained circulation in certain wild and farmed mammal species (10). Mammalian infections have frequently been associated with the acquisition of adaptive substitutions that enhance viral replication in non-avian hosts, with a broad spectrum of mammalian-adaptive mutations reported across the internal gene segments of avian influenza viruses (19, 20). Among these, the substitution at position 627 of the PB2 polymerase subunit, from glutamic acid to lysine (PB2-E627K), is one of the most widely observed and best characterised (23–26). PB2-E627K is regarded as a major determinant of increased viral replication, transcriptional activity, and pathogenicity in mammalian hosts. Consistent with this, PB2-E627K is more frequently detected in confirmed human H5N1 HPAIV infections and is therefore considered a key molecular marker in assessments of pre-pandemic potential (36). Mechanistically, this substitution enhances the compatibility of the viral polymerase complex with mammalian ANP32 proteins, thereby promoting efficient replication in mammalian cells (25, 28, 30). Despite conferring a clear replicative advantage in mammalian systems, PB2-627K remains rare among viruses isolated from avian hosts during the current panzootic. It remains unclear whether this reflects limited opportunities for spillback from mammals to birds or a fitness cost associated with PB2-627K in avian hosts.

In this study, we investigated the impact of the PB2-E627K substitution within a contemporary clade 2.3.4.4b genotype AB H5N1 HPAIV background, assessing the relative fitness of viruses carrying either PB2-627E or PB2-627K in two avian host species. We used ducks, representing aquatic waterfowl and the principal reservoir hosts for clade 2.3.4.4b H5N1 HPAIV, and chickens, as the most abundant farmed poultry species globally.

*In vitro* analyses demonstrated that PB2-E627K enhanced polymerase activity and supported increased replication in human cells, consistent with previous studies (22, 24, 26, 38–40). Importantly, this substitution did not reduce polymerase activity or viral replication in chicken or duck cells. These findings were corroborated by *in vivo* experiments in both species. Presence of PB2-627K did not dramatically affect infectivity, replication kinetics, shedding, pathological lesions, or tissue tropism, when present in a clade 2.3.4.4b H5N1 HPAIV compared with PB2-627E. Both variants productively infected all directly inoculated chickens and ducks, with vRNA consistently detected in both OP and C samples. vRNA was also detected in environmental samples, indicating comparable levels of environmental contamination by both variants. However, no statistically significant differences were observed in total vRNA shedding from either the OP or C in ducks or chickens infected with either PB2-627 variant. Small but nonsignificant differences in survival were observed between species and PB2-E627K containing isolates. Both PB2-627E and PB2-627K containing H5N1 HPAIVs transmitted efficiently between ducks, with 100% transmission. In contrast, transmission efficiency in chickens differed between variants (PB2-627K transmitted to 25% of R1s, whereas PB2-627E showed no transmission). Transmission of clade 2.3.4.4b HPAIV in chickens is frequently variable in experimental settings, and the small differences in transmission observed here may reflect stochastic processes rather than intrinsic biological differences. Nevertheless, previous studies have suggested that rapid mortality in chickens may limit opportunities for onward transmission (41). Consistent with this, chickens infected with H5N1 HPAIV containing PB2-627E died more rapidly than those infected with PB2-627K, potentially explaining the reduced transmission frequency observed. Conversely, ducks infected with PB2-627K exhibited a longer mean time to death and higher survival, yet transmission remained 100% efficient.

While relatively few studies have investigated the effect of PB2-627E or PB2-627K polymorphism in birds, as most tend to focus on consequences of mammalian adaptation, these findings are consistent with earlier studies of certain H5N1 gs/Gd lineages, including clade 2.2 viruses, in which PB2-627K replicated efficiently in avian cells (38). Similarly, no measurable fitness cost was observed in chickens infected with RG-derived isogenic clade 2.2 HPAIVs carrying either PB2-627E or PB2-627K (40). In other avian influenza subtypes, including H7N9 and H7N7, the presence of PB2-627K likewise did not negatively affect polymerase activity or viral replication fitness relative to PB2-627E (39, 42). In contrast, reduced fitness associated with PB2-627K has been reported in a pre-gs/Gd H5 HPAIV lineage, but not in later gs/Gd H5N1 HPAIV lineages such as clade 2.2 (38), supporting the notion that the phenotypic effects of this substitution are strongly dependent on viral genetic background. Within the contemporary clade 2.3.4.4b genotype AB background examined here, PB2-627K appears to be well tolerated in avian cells, with no detectable adverse impact on viral replication kinetics or polymerase activity.

A key finding of this study was the genetic stability of PB2-627K *in vivo*. Across all sequenced samples from directly infected and contact chickens and ducks, PB2-627K remained the consensus residue, with no evidence of reversion to PB2-627E. This contrasts with observations from some historic H5 HPAIV lineages in which PB2-627K was unstable in avian hosts (38). A small number of additional substitutions were detected, predominantly within polymerase genes recovered from ducks, but these were sporadic, were not associated with transmission chains, and showed no evidence of fixation. Collectively, these findings indicate that the contemporary clade 2.3.4.4b genotype AB polymerase complex is permissive to retention of PB2-627K. Nevertheless, this study evaluated only a single infection and transmission cycle under controlled experimental conditions, and additional ecological pressures encountered in wild bird populations may influence the long-term maintenance of this mutation.

Given the absence of a measurable fitness cost in chickens and ducks, the continued rarity of PB2-627K among avian detections is likely to reflect ecological and epidemiological constraints rather than negative selection within these hosts. Although mammalian infections have increased substantially during the clade 2.3.4.4b panzootic, most infections remain associated with predation or scavenging events and are often considered epidemiological dead ends (10, 43). Viral replication in these mammalian hosts is frequently concentrated within internal organs, reducing opportunities for environmental contamination and onward transmission relative to avian hosts (11, 16, 44, 45). Consequently, opportunities for successful transmission from infected mammals back into the wider avian reservoir may be uncommon. The low frequency of PB2-627K among avian viruses may therefore reflect limited opportunities for spillback rather than an inability of these viruses to circulate in birds.

Nevertheless, PB2-627K has been detected sporadically in avian-derived clade 2.3.4.4b viruses. For example, PB2-627K was identified in a white stork during the 2023 cat-associated outbreak in Poland (46). More strikingly however, surveillance data since 2022 have revealed a high prevalence of PB2-627K among European and American detections of clade 2.3.4.4b H5N5 (genotype I), particularly in seabirds and raptors (11, 47). While the disease ecology for H5N5 (genotype I) appears different to H5N1 clade 2.3.4.4b, and the reservoir species for H5N5 remains to be elucidated, these observations provide further evidence that PB2-627K can be stably maintained in avian populations within other genotypic backgrounds of clade 2.3.4.4b. However, the contrasting disease ecologies of clade 2.3.4.4b H5N1 and H5N5 subtypes suggest that host species and ecological context may modulate the fitness consequences of PB2-627K, with some avian hosts potentially tolerating, or even selecting for, this mammalian-adaptive substitution. In addition, experimental infection studies have demonstrated that certain avian taxa, particularly ratites, may support the emergence or maintenance of PB2-627K following infection due to differences in their ANP32 proteins (37, 48). Together with the findings presented here, these observations suggest that the consequences of PB2-627K are likely to depend on host species and viral genetic background. A more comprehensive understanding of ANP32 diversity across wild bird species may therefore improve identification of avian hosts permissive for the maintenance of mammalian-adaptive mutations (28–30).

Beyond clade 2.3.4.4b H5N1, evidence from other contemporary H5 HPAIV lineages further supports the capacity of mammalian-adaptive mutations to be maintained in avian hosts under certain genetic and ecological contexts. Notably, the PB2-627K polymorphism has been consistently detected in both human and avian infections with clade 2.3.2.1e (formerly 2.3.2.1c) H5N1 HPAIV in Cambodia since 2023 (49), indicating that this substitution can also be sustained in avian populations outside the clade 2.3.4.4b lineage. Interestingly, phylogenetic analyses of all eight gene segments revealed that H5N1 viruses detected in Cambodia from October 2023 onwards in both humans and poultry represent a novel reassortant genotype, generated through exchange between clades 2.3.2.1e and 2.3.4.4b, with the PB2 segment derived from clade 2.3.4.4b (49). However, whether selection for PB2-627K in this setting is driven by the PB2 segment alone or modulated by interactions with other viral gene segments remains unclear.

Importantly, not all mammalian-adaptive mutations appear to behave similarly in avian hosts. Recent work in yellow-legged gulls demonstrated that a mink-derived clade 2.3.4.4b H5N1 virus carrying multiple mammalian-associated substitutions, including PB2-T271A, exhibited markedly reduced infectivity and failed to transmit compared with an avian-derived counterpart (50, 51). Similarly, prolonged circulation in marine mammals has selected alternative polymerase adaptations, including PB2-701N and PB2-591K, highlighting that mammalian adaptation can follow multiple evolutionary pathways rather than converging on PB2-627K alone (12, 14, 44). Another residue of interest is PB2-627V, which has been reported in non-H5 influenza A viruses and appears to combine characteristics of avian-like PB2-627E and mammalian-adaptive PB2-627K (52). However, PB2-627V remains uncommon among clade 2.3.4.4b H5N1 viruses relative to PB2-627K (52), although it appears more frequently in H5N5 clade 2.3.4.4b, yet its phenotypic consequences in this lineage remain poorly understood.

These observations emphasise that mammalian adaptation should be considered a heterogeneous and context-dependent process, with the fitness consequences of individual mutations determined by both host species and viral genetic constellation. In contrast, the findings presented here indicate that PB2-627K is well tolerated within a contemporary clade 2.3.4.4b genotype AB background and does not impose a detectable fitness cost in chickens or ducks.

The present findings have important implications for avian influenza ecology, surveillance, and zoonotic risk assessment. We demonstrate that clade 2.3.4.4b H5N1 viruses carrying PB2-627K retain the ability to replicate efficiently, transmit, and remain genetically stable in two major avian host species without detectable reductions in fitness. These findings suggest that, if mammalian-adapted viruses are successfully reintroduced into avian populations, they may be capable of onward transmission and maintenance within those populations. Continued genomic surveillance in both avian and mammalian hosts is therefore important to detect spillback events and monitor the emergence and dissemination of viruses carrying established mammalian-adaptive markers associated with increased mammalian host competency.

## METHODS

### Viruses

Four AIV strains were used in this study, two field isolates of clade 2.3.4.4b H5N1 HPAIV, and two RG viruses. For *in vivo* investigations, two clade 2.3.4.4b H5N1 HPAIVs were used, both classified as Genotype AB (as designated by the European Union Reference Laboratory [EURL] GenIn tool (53); genotype B.18 as designated by the global genotype tool ggFlu (54)). The first virus, A/fox/England/017842/2022 (H5N1) ([GISAID EPI_ISL_18042582]), was isolated from a wild Red Fox (*Vulpes vulpes*) and possesses the PB2-627K mutation (hereafter referred to as ‘H5N1-627K’). The second virus, A/chicken/England/014330/2022 (H5N1) ([GISAID EPI_ISL_13370704]), was isolated from a domestic Chicken (*Gallus gallus domesticus*), and is the closest genetic relative to H5N1-627K across all UK poultry and wild bird AIV sequences generated to date, but which carried the PB2-627E residue (this strain is hereafter referred to as ‘H5N1-627E’). H5N1-627K and H5N1-627E were >99.47 % similar at the amino acid level across all genes (Table S3). For *in vitro* investigations, two RG viruses were also used, which were identical except in possessing either the PB2-627E or PB2-627K polymorphisms, these two viruses are hereby referred to as ‘RG-627E’ and ‘RG-627K’, respectively. Both viruses possessed the six internal gene segments (PB2, PB1, PA, NP, M and NS) from a genotype AB clade 2.3.4.4b H5N1 HPAIV (A/chicken/Scotland/054477/2021 [EPI_ISL_9012696]) as previously described (55). The remaining two gene segments, encoding the surface glycoproteins (HA and NA), were from the influenza A virus laboratory strain A/Puerto Rico/8/1934 (H1N1) (PR8). All viruses were propagated in specified pathogen-free (SPF) 9-day-old embryonated fowl eggs (EFEs) and titrated in EFEs to determine the 50% egg infectious dose (EID_50_), as previously described (41). The complete genomes of all virus stocks were sequenced, and minor variant analysis was performed at position 627 in PB2, to confirm that no amino acid polymorphisms had been introduced during egg passage relative to the original field isolates. For all *in vivo* infections, virus stocks were diluted in 0.1 M phosphate-buffered saline (PBS; pH 7.2).

### Ethics and Safety Statement

The *in vivo* experiments were reviewed and approved by the Animal and Plant Health Agency (APHA) Animal Welfare and Ethical Review Body to comply with the relevant UK legislation, in accord with the UK Home Office (HO) Project License PP7633638 and P9307748. According to the UK’s Advisory Committee on Dangerous Pathogens (ACDP) and the Specified Animal Pathogens Order (SAPO); wildtype gs/Gd lineage H5Nx HPAIVs are classified at ACDP hazard group level three pathogens and SAPO level four, hence all animal and laboratory work involving wildtype infectious material was conducted using licensed containment level 3 facilities at APHA. Work involving reverse genetics (RG) viruses was conducted in licensed containment facilities under an approved genetic modification (GM) risk assessment, in accordance with Health and Safety Executive (HSE) requirements and the guidance of the Scientific Advisory Committee on Genetic Modification (SACGM). Welfare monitoring of up to three times daily assessed the birds for humane endpoints following the onset of clinical disease, enabling any decisions to be made concerning the need for euthanasia. All birds had access to food and water *ad libitum*.

### Animals used and experimental design

High health status Pekin Ducks (*Anas platyrhynchos*) (Cherry Valley hybrid; Cherry Valley Farms Ltd., UK) were obtained at one-day old and reared until infection at 3 weeks of age. White Leghorn specified pathogen free (SPF) chickens (*Gallus gallus domesticus*) (Valo, Germany) were also reared at APHA from one day old until infection at 3 weeks of age. All birds were acclimatised for seven days prior to the first procedures which included swabbing (oropharyngeal (OP) and cloacal (C)) and bleeding from a superficial vein prior to inoculation. These samples were tested by M-gene real-time reverse transcription polymerase chain reaction (RT-PCR) (56) and antibody testing by hemagglutination inhibition (HI) (57) or ID Screen® Influenza A Antibody Competition Multi-species ELISA kit (IDVET, France), to exclude past AIV infection.

### *In vivo* duck and chicken experimental designs

Four donor (D0) ducks were inoculated via the ocular-nasal route with 100 µl per bird of either H5N1-627K or H5N1-627E at a titre of 10^4^ EID_50_. At 1 day post infection (dpi), eight recipient (R1) ducks were introduced to assess transmission dynamics. Similarly, six D0 chickens were inoculated via the ocular-nasal route with 100 µl per bird containing 10^5^ EID_50_ of either H5N1-627K or H5N1-627E, and eight R1 chickens were introduced at 1 dpi. Both OP and C swabs were collected daily from all birds between 1 and 14 dpi. Birds were humanely euthanised upon reaching predetermined clinical endpoints or at 14 dpi (study-end), when blood samples were collected via cardiac puncture under terminal anaesthesia. Opportunistic post-mortem examinations (PME) were conducted on ducks and chickens from both experimental groups when culled due to clinical endpoint decisions, and from culls of apparently healthy survivors at study-end. PMEs were performed on two ducks and two chickens infected with H5N1-627E, and on four ducks and three chickens infected with H5N1-627K. A comprehensive range of tissue samples was collected at PME for tissue tropism analyses.

Environmental samples, including drinking water and pond water from the ducks, along with drinking water, bedding straw, faeces, and air samples from the chickens, were collected on indicated days throughout the experiment and processed as previously described (41). Animals were classified as infected when vRNA was detected in either OP or C samples and this was accompanied by clinical disease and/or seroconversion.

### Clinical scoring and monitoring

All birds were monitored twice daily and assessed using a predefined clinical scoring system (Table S4). Following the onset of clinical signs, individual birds were subject to increased monitoring with an evening check to identify the humane endpoints. Birds were euthanised upon reaching a cumulative clinical score greater than or equal to 7 (Table S4). Any remaining individually housed birds were also euthanised, regardless of their clinical status.

### Clinical sample collection, processing and immunohistochemistry

Swabs were individually cut and placed into 1 ml of Leibovitz’s L-15 Medium containing antibiotics (LM; Gibco) as previously described (41). During PME, approximately 50 mg of tissue from selected organs was similarly collected into 1 ml LM and mechanically homogenised, with the clarified homogenate being used for RNA extraction. In addition, tissues were harvested from all major organs and fixed in 10% (v/v) neutral buffered formalin for a minimum period of 5 days before being embedded in paraffin. Four-micron thick serial sections were stained with haematoxylin and eosin (H&E), and for immunohistochemistry (IHC), using a mouse monoclonal anti-influenza A nucleoprotein antibody (Statens Serum Institute, Copenhagen, Denmark) as described (41). The overall distribution of virus-specific staining in each tissue was assessed using a semi-quantitative scoring system (where 0 = no staining, 1 = minimal, 2 = mild, 3 = moderate and 4 = widespread staining) (41). Specificity of immunolabelling was assessed in positive control sections by replacing the primary antibody with a matching mouse IgG isotype; no non-specific cross-linking was observed. Blood was collected prior to infection though wing bleeds, or at study end through cardiac puncture under terminal anaesthesia. Sera were separated from whole blood by incubation for 24 hours at 4 °C, followed by centrifugation for 5 minutes at 2000 rpm. Sera were heat inactivated at 56 °C for 30 minutes prior to serological analyses. Environmental samples were processed and total RNA extracted as described (41).

### RNA extraction and AIV reverse transcription Real-Time PCR (RT-PCR)

Total RNA was extracted from LM obtained from swab fluids, tissues, feathers, and environmental samples by using the MagMAX^TM^ CORE Nucleic Acid Purification Kit (Thermo Fisher Scientific™) as part of the robotic Mechanical Lysis Module (KingFisher Flex system; Life Technologies), according to the manufacturer’s instructions. Volumes of extracted RNA (2 µl) were tested by the M-gene (56) or H5 (58) specific RT-PCR using the primers and probes. RT-PCR Ct values <36 were considered as AIV positive, sub-threshold values in the range Ct 36.01-39.99 and Ct 40 (“No Ct”) were interpreted as negative. A ten-fold dilution series of titrated H5N1-627E and H5N1-627K viral RNA was used to construct a standard curve (Agilent, UK) to determine PCR efficiency, with Ct values obtained from viral specimens converted to relative equivalent units (REUs) by correlation with the EID_50_/ml values of the extracted viral standard on the standard curve as previously described (41). Birds were considered infected following positive detection of vRNA in either cavity, or by evidence of seroconversion obtained by HI or ELISA.

### Serological analysis

HI was conducted to determine the presence of antibodies against H5N1 subtypes according to standard methods (57). The duck sera were pre-absorbed by adding packed chicken red blood cells and incubated at 4°C overnight to prevent non-specific agglutination. Reciprocal HI titres of >1:16 were considered seropositive using internationally recognised criteria (57). All sera were also tested by the ID Screen® Influenza A Antibody Competition Multi-species ELISA (IDVET, France), which detects type-specific influenza A virus antibodies, targeted to the viral nucleoprotein (NP), according to the manufacturer’s instructions.

### Whole genome sequencing

Whole genome sequences were generated from positive samples using Oxford Nanopore Technology as described previously (59). Briefly, extracted vRNA was converted to double stranded cDNA and amplified by a one-step RT-PCR using SuperScript III One-Step RT-PCR kit (Thermo Fisher Scientific), see Table S5 for specific primers. PCR products were purified with Agencourt AMPure XP beads (Beckman Coultrer) prior to sequencing library preparation using the Native Barcoding Kit (Oxford Nanopore Technologies) and sequenced using a GridION Mk1 (Oxford Nanopore Technologies), according to manufacturer’s instructions (59). Assembly of the influenza A viral genomes were undertaken using a custom in-house pipeline described previously (17). To maximise sequencing success, samples selected had a RT-PCR Ct value <33. The python package flutile was used to identify consensus level amino acid polymorphisms in chosen samples using the H5N1-627K inocula as a reference (17).

### Genetic distribution of PB2 residue 627 variants across avian and mammalian families

All H5N1 clade 2.3.3.4b sequences were downloaded on 26/01/2026 from GISAID removing those currently under embargo. The dataset is available under EPI_SET ID EPI_SET_260409yp (https://doi.org/10.55876/gis8.260409yp). Duplicate sequences were removed using SeqKit v2.10.1. Sequences were subsequently filtered to exclude those lacking coverage at position 627, those derived from laboratory strains, or those for which host order could not be determined from the metadata or strain name. Minor populations such as A, D and V at position 627 were filtered for visualisation purposes but remain in the complete dataset (EPI_SET_260409). Sequences were aligned using MAFFT v7.505, manually trimmed to the open reading frame, and translated into amino acid sequences using AliView v2021. Amino acid frequencies at position 627 were calculated using Biopython v1.85 and pandas v2.2.3 and visualised using Matplotlib v3.10.0.

### Mini-replicon assays to measure influenza A virus polymerase activity

Expression plasmids for the EURL genotype AB H5N1 HPAIV (A/chicken/Scotland/054477/2021 [EPI_ISL_9012696]) minigenome assays were used as previously described (55). This virus naturally possesses the PB2-627E polymorphism, and this amino acid was mutated to the PB2-627K polymorphism using site directed mutagenesis, before being subcloned into pCAGGS as previously described (17, 55). Cells from Human (HEK 293T), chicken (CLEC213 and DF-1) or duck (CCL-141) (ATCC) were transfected in 24-well plates using lipofectamine 3000 transfection reagent (Thermo Fisher) with the following amount of pCAGGS-containing expression plasmids; 40ng of PB2, 40ng of PB1, 20ng of PA, 80ng of NP, 40ng of Renilla luciferase. Human cells were transfected with 80ng of the human pHOM1-Firefly plasmid and avian cells were transfected with 80ng of pCOM1-Firefly, as previously described (55). Cells were lysed 24 hours post-transfection using passive lysis buffer (Promega) and polymerase activity was measured using the Dual-Luciferase Reporter assay system (Promega) and a FLUOstar Omega plate reader (BMG Labtech). Firefly luciferase signal was normalised to Renilla luciferase signal to give relative luminescence units (RLU).

### Viral replication kinetics in avian cells

Confluent monolayers of primary chicken embryonic fibroblasts (CEFs) and duck embryonic fibroblasts (DEFs) were infected with 1 ml H5N1-627K or H5N1-627E multiplicity of infection (MOI) 0.001 in serum free DMEM (Gibco). After a 1-hour incubation at 37°C to allow viral attachment to cells, the inoculum was replaced with 2 ml of fresh virus growth media (DMEM, 0.15% BSA, 0.01% TPCK trypsin, 1% Pen/Strep). At 24, 48 and 72 hours post infection (hpi), supernatants were frozen, and samples stored at −80 °C until titrated by RT-PCR.

### Viral replication kinetics of RG viruses in avian and mammalian cells

Confluent monolayers of chicken lung epithelial cells (CLECs), duck embryonic fibroblasts (CCL-141s) and human alveolar basal epithelial cells (A549s) were infected with 200 µl of either RG-627E or RG-627K at an MOI of 0.001 in serum free DMEM (Gibco). After a 1-hour incubation at 37 °C, the inoculum was replaced with 2 mL of fresh virus growth media (DMEM, 0.01% TPCK trypsin). At 24, 48 and 72 hpi, supernatants were frozen, and samples stored at −80 °C until titrated on MDCK cells by plaque assay as previously described (60).

### Statistical analysis

Area under the curve (AUC) analysis of individual bird shedding profiles for material containing vRNA was undertaken using GraphPad Prism v10, with values expressed as REU as previously described (41). Birds that did not shed vRNA were assigned an AUC value of 1 REU. Mean AUC values were calculated and compared for statistical significance using one-way ANOVA with multiple comparisons in GraphPad Prism v10. Statistical analysis of viral growth kinetics data was performed using two-way ANOVA followed by Šídák’s multiple comparisons test using GraphPad Prism v10. Statistical significance was defined as p < 0.05. Kaplan–Meier survival curves were evaluated using the Gehan–Breslow–Wilcoxon test in GraphPad Prism v10.

## Acknowledgements

The authors would like to thank Nadia Chew, Amy Miller, Elena Mather, and Katie Brown for collection of samples from the *in vivo* experiments.

## Author Contributions

EB and SR contributed equally to this work. ACB and JJ contributed to Conceptualization. EB, SR, MDB, JQ, SJ, CDG, JY, JRS, ALS, KJR, JH, CJW, GW, DdS, AN, MJS and JJ contributed to Investigation. EB, SR, BM, ALS, AN, HC, BC and JJ contributed to Data curation. EB, SR, MDB, BM, SJ, JH, BC and HC contributed to Formal analysis. EB, SR, MDB, JQ, ACB and JJ contributed to Methodology. BM, BC and HC contributed to Software. EB, SR, BM and JJ contributed to Visualization. ACB and JJ contributed to Project administration and Resources. MI, WB, TP, ACB and JJ contributed to Supervision. EB, SR, SJ, ACB and JJ contributed to Writing – Original Draft. All authors contributed to Writing – Review & Editing. MI, WB, TP, ACB and JJ contributed to Funding acquisition. ACB and JJ jointly supervised this work.

## REFERENCES

1. Chen H. 2009. H5N1 avian influenza in China. Science in China Series C: Life Sciences 52:419–427.

2. Smith GJD, Donis RO, World Health Organization/World Organisation for Animal Health/Food Agriculture Organization H5 Evolution Working Group. 2015. Nomenclature updates resulting from the evolution of avian influenza A(H5) virus clades 2.1.3.2a, 2.2.1, and 2.3.4 during 2013–2014. Influenza and Other Respiratory Viruses 9:271–276.

3. Caliendo V, Lewis NS, Pohlmann A, Baillie SR, Banyard AC, Beer M, Brown IH, Fouchier RAM, Hansen RDE, Lameris TK, Lang AS, Laurendeau S, Lung O, Robertson G, van der Jeugd H, Alkie TN, Thorup K, van Toor ML, Waldenström J, Yason C, Kuiken T, Berhane Y. 2022. Transatlantic spread of highly pathogenic avian influenza H5N1 by wild birds from Europe to North America in 2021. Scientific Reports 12:11729.

4. Bevins SN, Shriner SA, Cumbee JC, Jr., Dilione KE, Douglass KE, Ellis JW, Killian ML, Torchetti MK, Lenoch JB. 2022. Intercontinental Movement of Highly Pathogenic Avian Influenza A(H5N1) Clade 2.3.4.4 Virus to the United States, 2021. Emerg Infect Dis 28:1006–1011.

5. Adlhoch C, Fusaro A, Gonzales JL, Kuiken T, Marangon S, Mirinaviciute G, Niqueux É, Stahl K, Staubach C, Terregino C, Broglia A, Baldinelli F. 2023. Avian influenza overview December 2022 – March 2023. EFSA Journal 21:e07917.

6. Banyard AC, Bennison A, Byrne AMP, Reid SM, Lynton-Jenkins JG, Mollett B, De Silva D, Peers-Dent J, Finlayson K, Hall R, Blockley F, Blyth M, Falchieri M, Fowler Z, Fitzcharles EM, Brown IH, James J. 2024. Detection and spread of high pathogenicity avian influenza virus H5N1 in the Antarctic Region. Nature Communications 15:7433.

7. Clessin A, Brusselmans M, Hong SL, Tornos J, Lejeune M, Shao Y, Briand FX, Abolnik C, Kaza B, Suchard MA, Aguado B, Alcamí A, Barbraud C, Beer M, Bennison A, Bonadonna F, Bonnet T, Bost CA, Boucheron S, Bralet T, Catry P, Cleeland J, Connan M, Coombes HA, Delord K, de Pasquale C, Dewar ML, Dong X, Emerit J, Fischer R, Fountain-Jones N, Galimberti F, González-Solís J, Guinet C, Gunn C, Günther A, Iervolino M, James J, Ji X, Jonsen I, Jones CW, Kuepfer A, Kuiken T, Lebohec C, Lisovski S, Llorente Zubiri L, Lynton-Jenkins JG, Martinez-García P, McCulley M, McMahon CR, et al. 2026. Dispersal, adaptation and persistence of H5N1 in the sub-Antarctic and Antarctica. bioRxiv doi:10.64898/2026.03.20.713283.

8. World Organisation for Animal Health (WOAH) World Animal Health Information System (WAHIS). 2026. Australia - Influenza A viruses of high pathogenicity (Inf. with) (non-poultry including wild birds) (2017-). https://wahis.woah.org/#/in-review/7649. Accessed July.

9. Knief U, Bouwhuis S, Globig A, Gunther A, Courtens W. 2026. Counting cases, conserving species: addressing highly pathogenic avian influenza in wildlife. Biol Rev Camb Philos Soc n/a:1464–7931.

10. Peacock TP, Moncla L, Dudas G, VanInsberghe D, Sukhova K, Lloyd-Smith JO, Worobey M, Lowen AC, Nelson MI. 2025. The global H5N1 influenza panzootic in mammals. Nature 637:304–313.

11. Falchieri M, Bentley E, Coombes HA, Mollett BC, Terrey J, Holland S, Stubbings E, Mcginn N, Cooper J, Ahmad S, Lewis J, Clifton B, Collinson N, Aegerter J, Venkatesh D, Russell DJF, James J, Reid SM, Banyard AC. 2025. Co-circulation of distinct high pathogenicity avian influenza virus (HPAIV) subtypes in a mass mortality event in wild seabirds and co-location with dead seals. bioRxiv doi:10.1101/2025.07.11.664278:2025.07.11.664278.

12. Pardo-Roa C, Nelson MI, Ariyama N, Aguayo C, Almonacid LI, Gonzalez-Reiche AS, Muñoz G, Ulloa M, Ávila C, Navarro C, Reyes R, Castillo-Torres PN, Mathieu C, Vergara R, González Á, González CG, Araya H, Castillo A, Torres JC, Covarrubias P, Bustos P, van Bakel H, Fernández J, Fasce RA, Johow M, Neira V, Medina RA. 2025. Cross-species and mammal-to-mammal transmission of clade 2.3.4.4b highly pathogenic avian influenza A/H5N1 with PB2 adaptations. Nature Communications 16:2232.

13. Plaza PI, Gamarra-Toledo V, Rodríguez Euguí J, Rosciano N, Lambertucci SA. 2024. Pacific and Atlantic sea lion mortality caused by highly pathogenic Avian Influenza A(H5N1) in South America. Travel Medicine and Infectious Disease 59:102712.

14. Uhart MM, Vanstreels RET, Nelson MI, Olivera V, Campagna J, Zavattieri V, Lemey P, Campagna C, Falabella V, Rimondi A. 2024. Epidemiological data of an influenza A/H5N1 outbreak in elephant seals in Argentina indicates mammal-to-mammal transmission. Nat Commun 15:9516.

15. Dhakal J, Bhat S, James J, Otwey RY, Chapagain S, Singh P. 2025. Highly Pathogenic Avian Influenza (HPAI) H5N1 in Raw Pet Foods and Milk: A Growing Threat to both Companion Animals and Human Health, and Potential Raw Pet Food Industry Liability. Journal of Food Protection 10.1016/j.jfp.2025.100628:100628.

16. Falchieri M, Reid SM, Dastderji A, Cracknell J, Warren CJ, Mollett BC, Peers-Dent J, Schlachter AD, McGinn N, Hepple R, Thomas S, Ridout S, Quayle J, Pizzi R, Núñez A, Byrne AMP, James J, Banyard AC. 2024. Rapid mortality in captive bush dogs (Speothos venaticus) caused by influenza A of avian origin (H5N1) at a wildlife collection in the United Kingdom. Emerg Microbes Infect 13:2361792.

17. Banyard AC, Coombes H, Terrey J, McGinn N, Seekings J, Clifton B, Mollett BC, Genova CD, Sainz-Dominguez P, Worsley L, Jorquera R, Billington E, Fullick E, Schlachter A-L, Jorge D, Núñez A, Falchieri M, James J, Reid SM. 2025. Detection of clade 2.3.4.4b H5N1 high pathogenicity avian influenza virus in a sheep in Great Britain, 2025. Emerging Microbes & Infections 14:2547730.

18. US Centers for Disease Control and Prevention (CDC). 2026. H5 Bird Flu: Current Situation. https://www.cdc.gov/bird-flu/situation-summary/index.html. Accessed 18th April 2026.

19. Long JS, Mistry B, Haslam SM, Barclay WS. 2019. Host and viral determinants of influenza A virus species specificity. Nature Reviews Microbiology 17:67–81.

20. Capelastegui F, Goldhill DH. 2025. H5N1 2.3.4.4b: a review of mammalian adaptations and risk of pandemic emergence. J Gen Virol 106.

21. Dholakia V, Quantrill JL, Richardson S, Pankaew N, Brown MD, Yang J, Capelastegui F, Masonou T, Case K-M, Ajeian J, Woodall MNJ, Magill C, Freimanis G, McCarron A, Staller E, Sheppard CM, Brown IH, Murcia PR, Smith CM, Iqbal M, Digard P, Barclay WS, Pinto RM, Peacock TP, Goldhill DH. 2025. Polymerase mutations underlie early adaptation of H5N1 influenza virus to dairy cattle and other mammals. bioRxiv doi:10.1101/2025.01.06.631435:2025.01.06.631435.

22. Mänz B, Schwemmle M, Brunotte L. 2013. Adaptation of avian influenza A virus polymerase in mammals to overcome the host species barrier. J Virol 87:7200–9.

23. MacCosham A, Vasiliu AG, Atchessi N. 2025. A rapid review of the avian influenza PB2 E627K mutation in human infection studies. Can Commun Dis Rep 51:137–144.

24. Subbarao EK, London W, Murphy BR. 1993. A single amino acid in the PB2 gene of influenza A virus is a determinant of host range. J Virol 67:1761–4.

25. Long JS, Idoko-Akoh A, Mistry B, Goldhill D, Staller E, Schreyer J, Ross C, Goodbourn S, Shelton H, Skinner MA, Sang H, McGrew MJ, Barclay W. 2019. Species specific differences in use of ANP32 proteins by influenza A virus. Elife 8.

26. Long JS, Giotis ES, Moncorgé O, Frise R, Mistry B, James J, Morisson M, Iqbal M, Vignal A, Skinner MA, Barclay WS. 2016. Species difference in ANP32A underlies influenza A virus polymerase host restriction. Nature 529:101–104.

27. Misra S, Gilbride E, Ramasamy S, Pond SLK, Kuchipudi SV. 2024. Enhanced Diversifying Selection on Polymerase Genes in H5N1 Clade 2.3.4.4b: A Key Driver of Altered Species Tropism and Host Range Expansion. bioRxiv doi:10.1101/2024.08.19.606826.

28. Carrique L, Fan H, Walker AP, Keown JR, Sharps J, Staller E, Barclay WS, Fodor E, Grimes JM. 2020. Host ANP32A mediates the assembly of the influenza virus replicase. Nature 587:638–643.

29. Domingues P, Hale BG. 2017. Functional Insights into ANP32A-Dependent Influenza A Virus Polymerase Host Restriction. Cell Rep 20:2538–2546.

30. Peacock Thomas P, Sheppard Carol M, Lister Margaret G, Staller E, Frise R, Swann Olivia C, Goldhill Daniel H, Long Jason S, Barclay Wendy S. 2023. Mammalian ANP32A and ANP32B Proteins Drive Differential Polymerase Adaptations in Avian Influenza Virus. Journal of Virology 97:e00213–23.

31. Staller E, Carrique L, Swann OC, Fan H, Keown JR, Sheppard CM, Barclay WS, Grimes JM, Fodor E. 2024. Structures of H5N1 influenza polymerase with ANP32B reveal mechanisms of genome replication and host adaptation. Nat Commun 15:4123.

32. Xie Z, Yang J, Jiao W, Li X, Iqbal M, Liao M, Dai M. 2025. Clade 2.3.4.4b highly pathogenic avian influenza H5N1 viruses: knowns, unknowns, and challenges. J Virol 99:e0042425.

33. Lindh E, Lounela H, Ikonen N, Kantala T, Savolainen-Kopra C, Kauppinen A, Österlund P, Kareinen L, Katz A, Nokireki T, Jalava J, London L, Pitkäpaasi M, Vuolle J, Punto-Luoma A-L, Kaarto R, Voutilainen L, Holopainen R, Kalin-Mänttäri L, Laaksonen T, Kiviranta H, Pennanen A, Helve O, Laamanen I, Melin M, Tammiranta N, Rimhanen-Finne R, Gadd T, Salminen M. 2023. Highly pathogenic avian influenza A(H5N1) virus infection on multiple fur farms in the South and Central Ostrobothnia regions of Finland, July 2023. Eurosurveillance 28:2300400.

34. Halwe NJ, Cool K, Breithaupt A, Schön J, Trujillo JD, Nooruzzaman M, Kwon T, Ahrens AK, Britzke T, McDowell CD, Piesche R, Singh G, Pinho dos Reis V, Kafle S, Pohlmann A, Gaudreault NN, Corleis B, Ferreyra FM, Carossino M, Balasuriya UBR, Hensley L, Morozov I, Covaleda LM, Diel DG, Ulrich L, Hoffmann D, Beer M, Richt JA. 2025. H5N1 clade 2.3.4.4b dynamics in experimentally infected calves and cows. Nature 637:903–912.

35. Kim DH, Lee DY, Seo Y, Song CS, Lee DH. 2025. Immediate PB2-E627K amino acid substitution after single infection of highly pathogenic avian influenza H5N1 clade 2.3.4.4b in mice. Virol J 22:183.

36. Briggs K, Kapczynski DR. 2023. Comparative analysis of PB2 residue 627E/K/V in H5 subtypes of avian influenza viruses isolated from birds and mammals. Front Vet Sci 10:1250952.

37. Coombes HA, Terrey J, Schlachter A-L, McCarter P, Regina I, Hepple R, McGinn N, Seekings J, Cooper J, Clifton B, Mollett BC, Falchieri M, Nunez A, Reid SM, James J, Banyard AC. 2025. Infection of ratites with clade 2.3.4.4b HPAIV H5N1: Potential implications for zoonotic risk. bioRxiv doi:10.1101/2025.09.08.674895:2025.09.08.674895.

38. Long JS, Howard WA, Núñez A, Moncorgé O, Lycett S, Banks J, Barclay WS. 2013. The effect of the PB2 mutation 627K on highly pathogenic H5N1 avian influenza virus is dependent on the virus lineage. J Virol 87:9983–96.

39. Gabriel G, Dauber B, Wolff T, Planz O, Klenk H-D, Stech J. 2005. The viral polymerase mediates adaptation of an avian influenza virus to a mammalian host. Proceedings of the National Academy of Sciences 102:18590–18595.

40. Bogs J, Kalthoff D, Veits J, Pavlova S, Schwemmle M, Mänz B, Mettenleiter TC, Stech J. 2011. Reversion of PB2-627E to −627K during replication of an H5N1 Clade 2.2 virus in mammalian hosts depends on the origin of the nucleoprotein. J Virol 85:10691–8.

41. James J, Billington E, Warren CJ, De Silva D, Di Genova C, Airey M, Meyer SM, Lewis T, Peers-Dent J, Thomas SS, Lofts A, Furman N, Nunez A, Slomka MJ, Brown IH, Banyard AC. 2023. Clade 2.3.4.4b H5N1 high pathogenicity avian influenza virus (HPAIV) from the 2021/22 epizootic is highly duck adapted and poorly adapted to chickens. Journal of General Virology 104.

42. Mok Chris Ka P, Lee Horace Hok Y, Lestra M, Nicholls John M, Chan Michael Chi W, Sia Sin F, Zhu H, Poon Leo Lit M, Guan Y, Peiris Joseph Sriyal M. 2014. Amino Acid Substitutions in Polymerase Basic Protein 2 Gene Contribute to the Pathogenicity of the Novel A/H7N9 Influenza Virus in Mammalian Hosts. Journal of Virology 88:3568–3576.

43. Plaza PI, Gamarra-Toledo V, Euguí JR, Lambertucci SA. 2024. Recent Changes in Patterns of Mammal Infection with Highly Pathogenic Avian Influenza A(H5N1) Virus Worldwide. Emerg Infect Dis 30:444–452.

44. Banyard A, Mollett B, Lynton-Jenkins J, Richardson S, Quantrill J, Byrne A, Harvey R, Adams L, Proust A, Brown M, Yang J, Reid S, Coombes H, Clifton B, Bennison A, Clessin A, Gamble A, Fitzcharles E, Fowler Z, Masonou T, Case K, Aejian J, Woodall M, Smith C, Iqbal M, Brown I, Lewis N, Barclay W, Peacock T, James J. 2025. Disease ecology and zoonotic risk of clade 2.3.4.4b H5N1 high pathogenicity avian influenza in the sub-Antarctic region. Research Square [Preprint] doi:10.21203/rs.3.rs-8029950/v1.

45. Castillo A, Fasce R, Parra B, Andrade W, Covarrubias P, Hueche A, Campano C, Tambley C, Rojas M, Araya M, Hernández F, Bustos P, Fernández J. 2023. The first case of human infection with H5N1 avian Influenza A virus in Chile. J Travel Med 30.

46. Rabalski L, Milewska A, Pohlmann A, Gackowska K, Lepionka T, Szczepaniak K, Swiatalska A, Sieminska I, Arent Z, Beer M, Koopmans M, Grzybek M, Pyrc K. 2023. Emergence and potential transmission route of avian influenza A (H5N1) virus in domestic cats in Poland, June 2023. Euro Surveill 28.

47. Erdelyan CNG, Kandeil A, Signore AV, Jones MEB, Vogel P, Andreev K, Bøe CA, Gjerset B, Alkie TN, Yason C, Hisanaga T, Sullivan D, Lung O, Bourque L, Ayilara I, Pama L, Jeevan T, Franks J, Jones JC, Seiler JP, Miller L, Mubareka S, Webby RJ, Berhane Y. 2024. Multiple transatlantic incursions of highly pathogenic avian influenza clade 2.3.4.4b A(H5N5) virus into North America and spillover to mammals. Cell Rep 43:114479.

48. Baker SF, Ledwith MP, Mehle A. 2018. Differential Splicing of ANP32A in Birds Alters Its Ability to Stimulate RNA Synthesis by Restricted Influenza Polymerase. Cell Rep 24:2581–2588.e4.

49. Siegers JY, Xie R, Byrne AMP, Edwards KM, Hu S, Yann S, Sin S, Tok S, Chea K, Horm S, Rith C, Keo S, Pum L, Duong V, Auerswald H, Phou Y, Kol S, Spiegel A, Harvey R, Tum S, Sorn S, Seng B, Sengdoeurn Y, Darapheak C, Savuth C, Hak M, Ieng V, Patel S, Thielen P, Claes FF, Lewis NS, Sovann L, Dhanasekaran V, Karlsson EA. 2025. Emergence of a Novel Reassortant Clade 2.3.2.1c Avian Influenza A/H5N1 Virus Associated with Human Cases in Cambodia. medRxiv doi:10.1101/2024.11.04.24313747:2024.11.04.24313747.

50. Agüero M, Monne I, Sánchez A, Zecchin B, Fusaro A, Ruano MJ, Del Valle Arrojo M, Fernández-Antonio R, Souto AM, Tordable P, Cañás J, Bonfante F, Giussani E, Terregino C, Orejas JJ. 2023. Highly pathogenic avian influenza A(H5N1) virus infection in farmed minks, Spain, October 2022. Euro Surveill 28.

51. Puente AP, Tarrés-Freixas F, Bessière P, Gaide N, Martín L, Contreras E, Martorell J, Nofrarías M, Valle R, Girbau A, Pérez M, Pérez M, Valdez-May MJ, Reska T, Cordón I, Majó N, Guérin J-L, Urban L, Bertran K. 2025. From feather to fur: gull and mink H5N1 clade 2.3.4.4b HPAIV in their original hosts and their spillover and spillback potential. Research Square [Preprint] doi:10.21203/rs.3.rs-7357485/v1.

52. Guo Y, Shu S, Zhou Y, Peng W, Jiang Z, Li Y, Li T, Du F, Wang L, Chen X, Dong J, Zhao C, Haitian Wang M, Sun Y, Sun H, Lu L, Digard P, Chang KC, Yen HL, Liu J, Pu J. 2025. An emerging PB2-627 polymorphism increases the zoonotic risk of avian influenza virus by overcoming ANP32 host restriction in mammalian and avian hosts. J Virol 99:e0085325.

53. (IZSVe) IZSdV. 2025. Genin2 (v2.1.6). https://github.com/izsvenezie-virology/genin2. Accessed 20th July 2026.

54. ggFlu. 2026. ggFlu: Tools for Global Genotyping of Influenza Virus. https://www.ggflu.com/. Accessed 20 July 2026.

55. Harvey WT, Pinto RM, Brown MD, Lu L, Quantrill JL, Yang J, Pankaew N, Nel M, Baxter J, Byrne AMP, Kapczynski DR, Iqbal M, James J, Banyard AC, Brown I, Barclay W, Peacock TP, Digard P, Lycett SJ. 2025. Genetic reassortment and diversification of host specificity have driven evolutionary trajectories of lineages of panzootic H5N1 influenza. bioRxiv doi:10.1101/2025.08.20.670882:2025.08.20.670882.

56. Nagy A, Černíková L, Kunteová K, Dirbáková Z, Thomas SS, Slomka MJ, Dán Á, Varga T, Máté M, Jiřincová H, Brown IH. 2021. A universal RT-qPCR assay for “One Health” detection of influenza A viruses. PLOS ONE 16:e0244669.

57. World Organisation for Animal Health (WOAH). 2021. Terrestrial Manual: Chapter 3.3.4 Avian Influenza (including infection with high pathogenicity avian influenza viruses).

58. James J, Seekings AH, Skinner P, Purchase K, Mahmood S, Brown IH, Hansen RDE, Banyard AC, Reid SM. 2022. Rapid and sensitive detection of high pathogenicity Eurasian clade 2.3.4.4b avian influenza viruses in wild birds and poultry. J Virol Methods 301:114454.

59. Animal and Plant Health Agency (APHA). 2026. Next Generation Sequencing using Oxford Nanopore Technology for Influenza Viruses and Notifiable Avian Disease Genotyping and Pathotyping, *on* FluGlobalNet. https://science.vla.gov.uk/fluglobalnet/Documents/english/protocol_ONT.pdf. Accessed 23 Feb 2026.

60. James J, Bhat S, Walsh Sarah K, Karunarathna Thusitha K, Sadeyen J-R, Chang P, Sealy Joshua E, Mahmood S, Mollett Benjamin C, Slomka Marek J, Brookes Sharon M, Iqbal M. 2022. The Origin of Internal Genes Contributes to the Replication and Transmission Fitness of H7N9 Avian Influenza Virus. Journal of Virology 96:e01290–22.

