## Supplementary Figures for "The mammalian-adaptive PB2-E627K substitution preserves viral fitness of clade 2.3.4.4b H5N1 HPAIV in birds"

### Figure S1

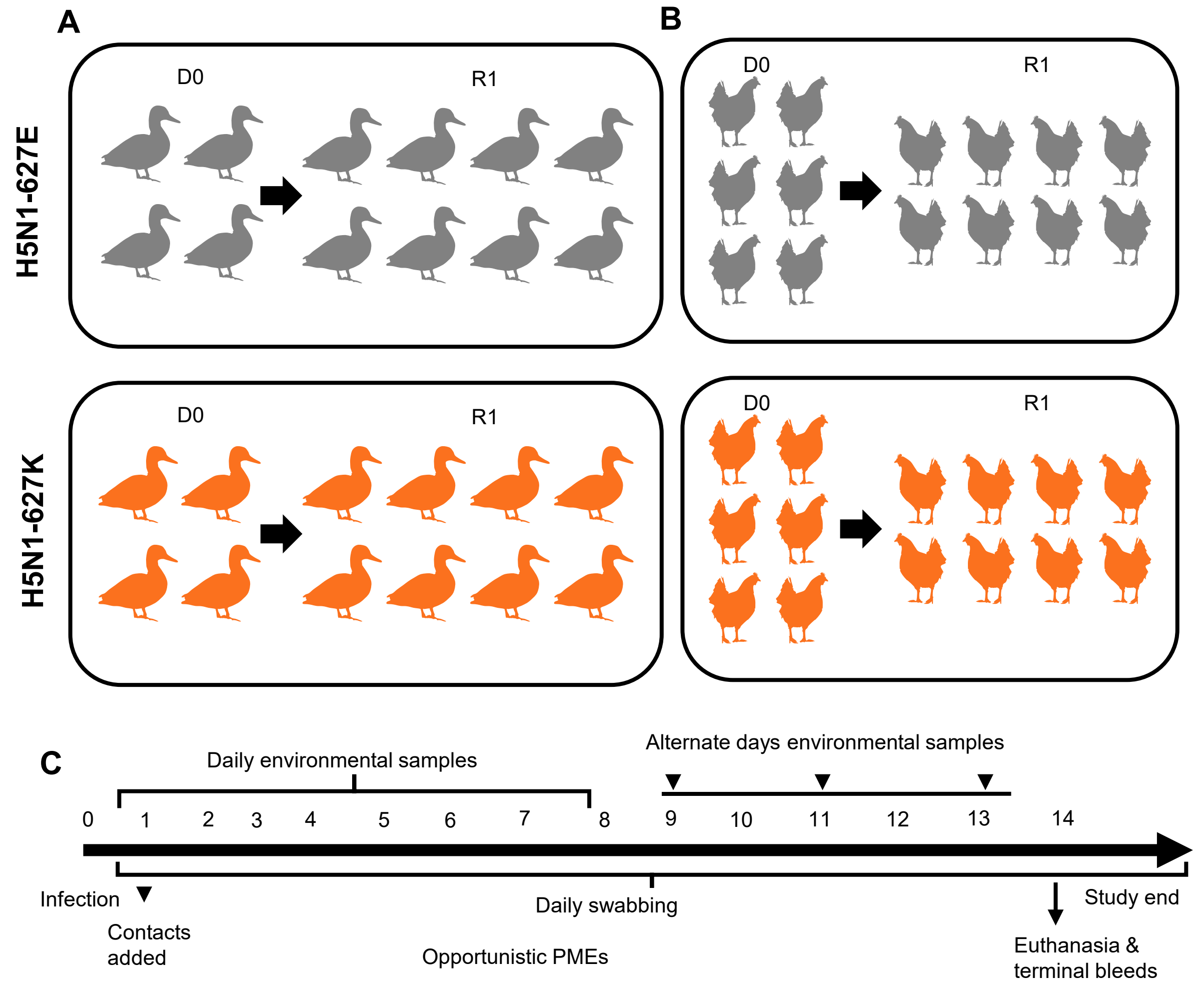


D

| Virus | Species | D0 bird ID | R1 bird ID |
|---|---|---|---|
| H5N1-627E | Chicken | 3515-3520 (n = 6) | 3521-3528 (n = 8) |
| H5N1-627E | Duck | 236-239 (n = 4) | 240-247 (n = 8) |
| H5N1-627K | Chicken | 3529-3534 (n = 6) | 3535-3542 (n = 8) |
| H5N1-627K | Duck | 125-128 (n = 4) | 129-136 (n = 8) |

**Figure S1. Schematic of the in vivo experimental design to assess infection, replication, transmission, and survival kinetics in ducks and chickens.**
Groups of ducks (**A**; n=4) and chickens (**B**; n=6) were inoculated (D0) via the ocular–nasal route with 100 µl per bird of $10^5$ $EID^{50}$ of Clade 2.3.4.4b H5N1 viruses, which were genetically highly similar, yet differed in the amino acid at position 627 in the PB2 protein, either 627E (H5N1-627E) or K H5N1-627K). At 1 day post-infection (dpi), contact birds (R1; n=8 per species) were introduced to assess transmission. **C**; Oropharyngeal (OP) and cloacal (C) swabs were collected daily from all birds between 1–14 dpi. Birds were humanely euthanised upon reaching predefined clinical endpoints or at 14 dpi, when blood samples were obtained via cardiac puncture under terminal anaesthesia. Opportunistic post-mortem examinations (PMEs) were conducted on birds euthanised due to clinical signs or at study termination. Environmental samples were collected daily from 1–8 dpi and on alternate days from 9–14 dpi. **D**; Bird numbers in each group.

### Figure S2

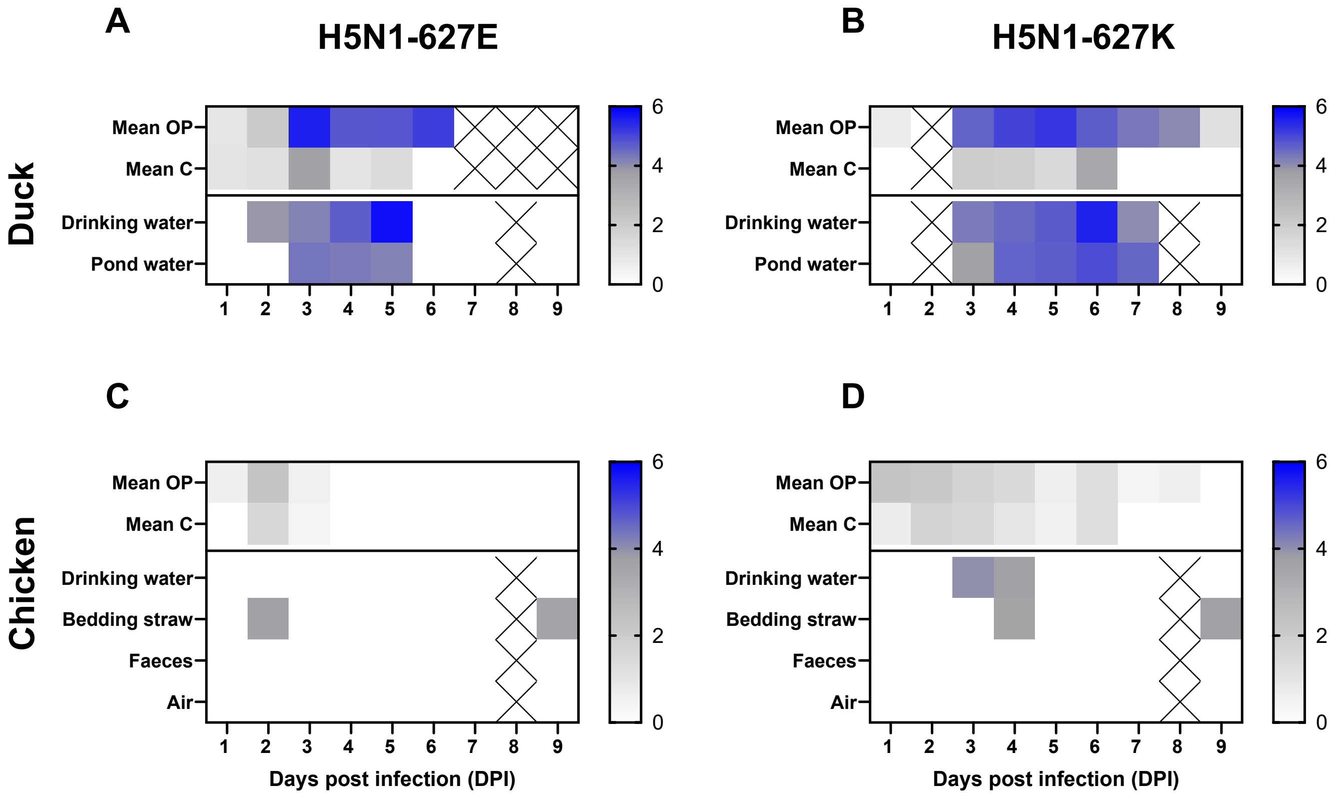


**Figure S2. Environmental contamination associated with infection and transmission of clade 2.3.4.4b H5N1 HPAIVs carrying PB2-627E or PB2-627K.**
Environmental viral RNA (vRNA) levels are shown as relative equivalency units (REU) based on RT-PCR analysis of RNA extracted from samples collected from the environments of ducks (A and B) and chickens (C and D). Birds were inoculated with clade 2.3.4.4b H5N1 viruses that were genetically near-identical, differing at position 627 of the PB2 protein, either E (H5N1-627E; A and C) or K (H5N1-627K; B and D). Contact ducks and chickens were introduced at 1-day post-infection (dpi) to assess transmission. Colour intensity reflects the quantity of vRNA detected, with blue shades indicating higher levels, grey showing >Ct 36, and white indicating undetectable vRNA. “X” symbols denote samples that were either not collected or could not be tested.

### Figure S3

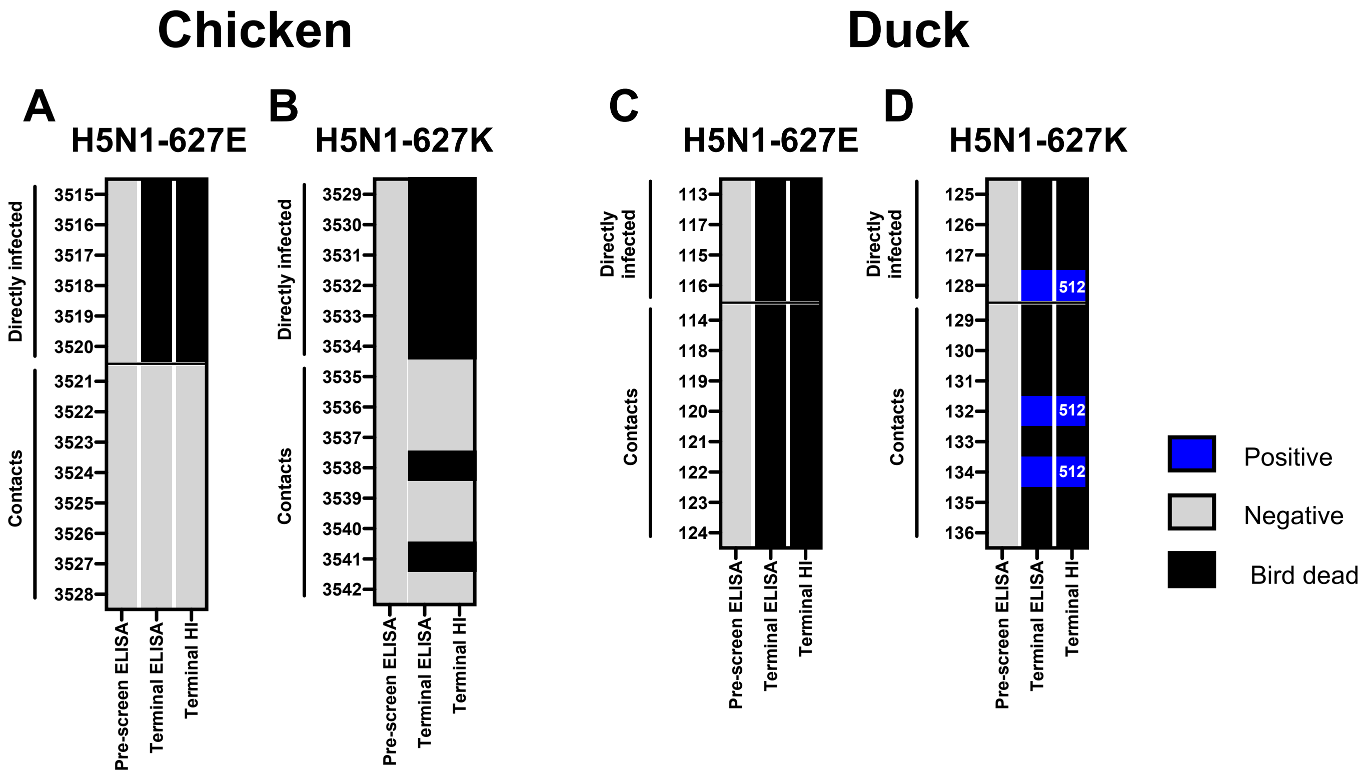


**Figure S3. Seroconversion of chickens and ducks following infection or contact exposure with clade 2.3.4.4b H5N1 HPAIVs carrying PB2-627E or PB2-627K.**

Serum samples were collected from chickens (**A** and **B**) and ducks (**C** and **D**) that were either directly inoculated with H5N1-627K or H5N1-627E viruses, or co-housed with inoculated birds as contact-exposed animals. Seroconversion was assessed using a haemagglutination inhibition (HI) assay with a homologous clade 2.3.4.4b H5N1 HPAIV antigen. HI titres are shown for individual birds that survived to 14 days post-infection (dpi) (terminal). Sera collected prior to challenge and at 14 dpi (terminal) were also tested by ELISA for antibodies against the influenza A virus nucleoprotein (NP). Negative results are shown in grey, positive results in blue, and birds that died before the terminal sampling point (and therefore had no serum sample available) are shown in black.
