## Supplementary Tables for "The mammalian-adaptive PB2-E627K substitution preserves viral fitness of clade 2.3.4.4b H5N1 HPAIV in birds"

**Table S1.** Summary of infection and transmission in chickens and ducks

| Species | Virus | Directly infected birds | | | Contact birds | |
|---|---|---|---|---|---|---|
| | | Infection (%) | Mortality (%) | Mean Death Time (days) | Transmission (%) | Mean day post contact to first vRNA positive (SD) |
| Chicken | H5N1-627E | 100 | 100 | 2.17 | 0 | n/a |
| Duck | | 100 | 100 | 4.5 | 100 | 1.88 (±0.35) |
| Chicken | H5N1-627K | 100 | 100 | 3.3 | 25 | 3.5 (±0.71) |
| Duck | | 100 | 75 | 6 | 100 | 2.25 (±0.46) |

SD = standard deviation

**Supplementary Tables**

**Table S2.** Genetic polymorphisms identified in clade 2.3.4.4b H5N1 HPAIV following infection and transmission in chickens and ducks.

| **Segment** | **Position** | **Ref** | **Variant** | **Sample ID** | **Tissue/Sample** | **Timepoint** | **Species** | **D0/R1** |
|---|---|---|---|---|---|---|---|---|
| PB2 | 553 | V | I | 134 | Brain | PM 14 dpi | Duck | R1 |
| PB2 | 117 | T | P | 135 | OP swab | 3 dpi | Duck | R1 |
| PB2 | 718 | K | R | 128 | OP swab | 6 dpi | Duck | R1 |
| PB2 | 62 | R | G | 136 | Heart | PM 5.2 dpi | Duck | D0 |
| PB2 | 666 | T | Q | 3531 | Muscle | PM 2 dpi | Chicken | R1 |
| PB2 | 672 | A | G | 130 | OP swab | 5 dpi | Duck | D0 |
| PB2 | 673 | G | D | 130 | OP swab | 5 dpi | Duck | R1 |
| PB1 | 709 | V | I | 3531 | C swab, proventriculus, muscle, pancreas, lung, liver, kidney, intestine, heart, caecal tonsil | 2 dpi and PM 2 dpi | Chicken | D0 |
| PB1 | 9 | F | S | 134 | OP swab | 6 dpi | Duck | R1 |
| PA | 357 | T | A | 134 | OP swab | 6 dpi | Duck | R1 |
| NP | 375 | D | A | 128 | OP swab | 6 dpi | Duck | D0 |
| NA | 44 | Y | H | 134 | Brain | PM 14 dpi | Duck | R1 |
| NS | 142 | E | V | 134 | Brain | PM 14 dpi | Duck | R1 |

HA and M have no substitutions reported. Ref = reference sequence of the inoculum. Amino acid positions were numbered from the N-terminal methionine of each influenza protein of A/fox/England/017842/2022 (H5N1). D0 = directly infected, R1 = contact. DPI = days post infection, PM = post mortem.

### Supplementary Tables

**Table S3.** Sequence identity between H5N1-627E and H5N1-627K

| Gene Segment | Sequence similarity (%) |
|---|---|
| PB2 | 99.47 |
| PB1 | 100 |
| PA | 99.86 |
| HA | 99.82 |
| NP | 100 |
| NA | 99.57 |
| MP | 100 |
| NS | 99.64 |

**Supplementary Tables**

**Table S4.** Total counts of each clinical sign observed following infection with H5N1 AB-mammalian and AB-avian

| Clinical Sign | Clinical sign severity score | Clinical Score Incidence Totals | | | | | | | |
|---|---|---|---|---|---|---|---|---|---|
| | | H5N1-627-K | | | | H5N1-627-E | | | |
| | | Chicken | | Duck | | Chicken | | Duck | |
| | | D0 | R1 | D0 | R1 | D0 | R1 | D0 | R1 |
| **Changes in huddling** | **1** | 11 | 3 | 2 | 1 | 7 | 0 | 3 | 4 |
| **Eyes closed** | **1** | 9 | 1 | 3 | 0 | 2 | 0 | 5 | 2 |
| **Conjunctivitis** | **1 - 2** | 0 | 0 | 6 | 15 | 1 | 0 | 0 | 3 |
| **Changes in body position** | **1** | 17 | 3 | 3 | 5 | 6 | 0 | 7 | 11 |
| **Oedema** | **2** | 1 | 0 | 8 | 0 | 3 | 0 | 2 | 1 |
| **Cyanosis of extremities** | **2-3** | 1 | 0 | 0 | 3 | 0 | 0 | 0 | 0 |
| **Lethargy / reluctance to fly*** | **1** | 17 | 3 | 6 | 13 | 8 | 0 | 9 | 38 |
| **Lack of engagement with enrichment** | **1** | 2 | 1 | 1 | 4 | 2 | 0 | 2 | 16 |
| **Oronasal discharge** | **2** | 2 | 0 | 1 | 1 | 0 | 0 | 0 | 0 |
| **Diarrhoea** | **2-3** | 4 | 1 | 0 | 0 | 5 | 1 | 1 | 5 |
| **Perceived weight reduction** | **2** | 0 | 0 | 1 | 1 | 0 | 0 | 0 | 0 |
| **Loss of balance** | **2** | 5 | 1 | 1 | 2 | 5 | 0 | 0 | 0 |
| **Tremors** | **2 - 3** | 4 | 1 | 2 | 3 | 4 | 0 | 2 | 3 |
| **Torticollis** | **4** | 0 | 0 | 0 | 0 | 0 | 0 | 0 | 0 |
| **Seizure#** | **3, 4, 7** | 0 | 0 | 1 | 5 | 0 | 0 | 5 | 7 |
| **Paralysis/inability to eat or drink** | **7** | 1 | 0 | 2 | 0 | 1 | 0 | 1 | 0 |
| **Found dead** | **N/A** | 1 | 1 | 0 | 0 | 1 | 0 | 0 | 0 |
| **Euthanised** | **N/A** | 5 | 1 | 3 | 6 | 5 | 0 | 4 | 7 |
| **Total instances** | | **80** | **16** | **40** | **59** | **50** | **1** | **41** | **97** |

Range in score depends on severity of the clinical sign. *scored for chickens only; # scored for ducks only. D0 = directly infected, R1 = contact.

### Supplementary Tables

**Table S5.** Primers used throughout the study

| Primer | Sequence 5'-3' | Application |
|---|---|---|
| Optil-F1 | TTACGCGCCAGCAAAAGCAG | RT-PCR for ONT |
| Optil-F2 | GTTACGCGCCAGCGAAAGCAGG | RT-PCR for ONT |
| Optil-R1 | GTTACGCGCCAGTAGAAACAAG | RT-PCR for ONT |
